# The RALF Signaling Pathway Regulates Cell Wall Integrity during Pollen Tube Growth in Maize

**DOI:** 10.1101/2023.01.24.525389

**Authors:** Liang-Zi Zhou, Lele Wang, Zengxiang Ge, Julia Mergner, Xingli Li, Bernhard Küster, Gernot Längst, Li-Jia Qu, Thomas Dresselhaus

## Abstract

Autocrine signaling pathways regulated by RAPID ALKALINIZATION FACTORs (RALFs) control cell wall integrity during pollen tube germination and growth in Arabidopsis. To investigate the role of pollen-specific RALFs in another plant species, we combined gene expression data with phylogenetic and biochemical studies to identify candidate orthologs in maize. We show that Clade IB *ZmRALF2/3* mutants, but not that of Clade III *ZmRALF1/5* caused cell wall instability in the sub-apical region of the growing pollen tube. ZmRALF2/3 are mainly located to the cell wall and are partially able to complement the pollen germination defect of their Arabidopsis orthologs AtRALF4/19. Mutations in *ZmRALF2/3* compromise pectin distribution pattern leading to altered cell wall thickness, hyperphosphorylation of ZmPEX cell wall proteins and pollen tube burst. Clade IB, but not Clade III ZmRALFs are capable to interact with pollen-specific CrRLK1L receptor kinases ZmFERL4/7/9 and GPI-anchored co-receptors ZmLLG1/2 at similar binding affinities. In contrast, binding affinity to ZmPEX2/4 cell wall proteins is about five times higher. Based on these data, we now propose a dosage-dependent model showing how Clade IB RALFs act as extracellular sensors to regulate cell wall integrity and thickness during pollen tube growth in plants.

**One sentence summary:** Pollen-specific RALFs interact at different binding affinities with receptor kinases, GPI-anchored proteins and cell wall proteins to regulate cell wall integrity during pollen tube growth in maize.

## INTRODUCTION

Flowering plants (angiosperms) have evolved pollen tubes that carry their passive and immobile sperm cell cargo deep into the maternal reproductive tissues towards the embryo sac for double fertilization (Dresselhaus et al., 2016). During their journey from the stigma through the transmitting tract of the style towards ovules, pollen tubes secrete many small proteins for communication with various maternal tissues (Ge et al., 2019a; Johnson et al., 2019; Kim et al., 2021). These include RAPID ALKALINIZATION FACTORs (RALFs) that were shown to regulate hydration as well as cell wall integrity during pollen germination, tube growth and reception in the model plant Arabidopsis (Ge et al., 2017; Mecchia et al., 2017; Liu et al., 2021; Gao et al., 2022; Zhong et al., 2022). First discovered in tobacco leaves (Pearce et al., 2001), *RALF* genes were found to be universally distributed among land plants with one to three family members in mosses and up to 37 members in eudicots like Arabidopsis (Campbell and Turner, 2017; Abarca et al., 2021). Functional studies have been mainly performed in Arabidopsis: it was shown, for example, that *AtRALF1* overexpression or application of the peptide inhibited rapid root growth (Matos et al., 2008; Li et al., 2022), while downregulation showed increased root length (Bergonci et al., 2014). AtRALF34 was reported as part of a signaling network that regulates lateral root initiation (Gonneau et al., 2018). Leaf expressed AtRALF23 inhibits immune responses (Stegmann et al., 2017; Gronnier et al., 2022) and induces stomatal closure (Yu et al., 2018). *RALFs* were also reported to play important roles in reproduction: initially it was shown in tomato that *in vitro* application of pollen-specific SlPRALF inhibited pollen germination and it interacts with pollen-specific leucine-rich repeat (LRR)/extensin-like chimera proteins (LRXs) (Covey et al., 2010). In Arabidopsis, it was later shown that AtRALF23/33 regulate hydration of pollen grains at the stigma (Liu et al., 2021), while AtRALF4/19 redundantly regulate pollen germination and tube growth. It was shown that this regulation occurs via binding to LRX cell wall proteins (Mecchia et al., 2017; Wang et al., 2018), as well as an autocrine signaling pathway via binding to pollen-specific *Catharanthus roseus* receptor-like kinase 1-like (CrRLK1Ls) family members ANXUR 1 (ANX1), ANX2, BUDDA’s PAPER SEAL 1 (BUPS1) and BUPS2 (Ge et al., 2017). It was further reported that GPI-anchored LORELEI-like proteins LLG2 and LLG3 serve as co-receptors in BUPS/ANX-RALF signaling to regulate integrity during pollen tube germination and growth in Arabidopsis (Feng et al., 2019; Ge et al., 2019c). Recently, AtRALF6/7/16/36/37 were shown to establish a polytubey block and to control pollen tube reception and rupture inside targeted embryo sacs via CrRLK1L family members FERONIA (FER)1, ANJEA (ANJ) and HERCULES RECEPTOR KINASE 1 (HERK1) (Zhong et al., 2022). Notably, another report suggests that AtRALF4/19 act via the FER-LRE module to induce pollen tube rupture and sperm release (Gao et al., 2022).

Structural studies elucidated the interactions showing using AtRALF23 as an example that binds to a surface groove of LLG2 with a conserved N-terminal recognition motif and thereby mediates interaction with FER forming a heterotrimeric complex (Xiao et al., 2019). Crystal structures of LRX2 and LRX8 binding with AtRALF4 revealed a dimeric arrangement of LRX proteins of which each monomer directly interacts with its N-terminal LRR domain to one RALF peptide (Moussu et al., 2020).

Compared with Arabidopsis, little is known about RALF-LLG-CrRLK1L and RALF-LRX signaling pathways in other plant species. With the exception of the LRX proteins PEX1/2 (Rubinstein et al., 1995b; Rubinstein et al., 1995a) that were reported to affect pollen tube growth in maize (*Zea mays*), nothing has been reported to our knowledge about other components of these pathways during reproduction in grasses like maize, a species that probably generates the fastest growing pollen tubes. Pollen hydration and germination occurs in maize within 5 minutes after landing on papilla silk hair structures (Bedinger, 1992), before pollen tubes penetrate the silk and enter the transmitting tract where they grow up to 30 cm at a speed of more than 10 mm per hour towards the surface of the ovule. They are further guided towards the micropylar region of the ovule, grow inside and release its sperm cell cargo into the receptive synergid cell (Zhou et al., 2017). It was speculated that maize pollen tubes secrete various small proteins along their journey to trigger, for example, papilla hair cell recognition and penetration, nutrient release and cell wall softening during transmitting tract growth, ovule penetration as well as synergid cell death and pollen tube rupture (Dresselhaus et al., 2011).

In order to study the role of pollen-specific *RALF* genes in maize, we searched the genome of the inbred line B73 for members of these gene families and combined expression and phylogenetic studies with biochemical interaction studies to elucidate similarities and differences to above reported findings in Arabidopsis. Three RALF clades were identified and pollen-expressed Clade IB and Clade III ZmRALFs were compared throughout this report. *ZmRALFs* levels were down-regulated by RNAi to investigate milder phenotypes as well as fully knocked out by CRISPR/cas9. Pollen germination and tube growth as well as cell wall composition and thickness was analyzed in the mutants. Binding affinities were compared between ZmRALF interaction partners and a phosphoproteomics approach was conducted to investigate consequences of reduced ZmRALFs signaling. Finally, we discuss a model how the RALF interaction partners are interconnected and propose Clade IB ZmRALFs as sensors of cell wall integrity, composition and thickness during pollen tube growth.

## RESULTS

### Pollen-Expressed Maize RALFs Belong to Clade IB or Clade III

RNA-seq data was generated from pollen grains, pollen tubes, other reproductive and vegetative tissues of the maize inbred line B73 and deposited in the CoNekT online database (https://evorepro.sbs.ntu.edu.sg). Among the most strongly and specifically expressed genes in pollen grains and pollen tubes, we identified three genes encoding *RALFs* with TPM (transcript per million) expression values >7,000 (Supplemental Figure S1). Protein sequences of these three genes named as *ZmRALF1/2/3* were used as a query for various genome wide BLAST searches to identify all members of the *RALF* gene family in maize. In total, 24 *RALF*-like genes were identified of which nine are expressed in pollen grains and tubes. *ZmRALF1/2/3/5* each show a strong (>2,500 TPM) and pollen-specific expression pattern, while *ZmRALF4/6/7/8* each show moderate (50-400 TPM) expression levels. *ZmRALF9* is lowly expressed in pollen, but strongly expressed in roots. Other members like *ZmRALF12* are broadly expressed in vegetative tissues like stigma, root, leaf, and seed (Supplemental Figure S1).

The RALF family in Arabidopsis is expanded and contains 37 members (Campbell and Turner, 2017; Abarca et al., 2021). Phylogenetic analysis was done by using and comparing the 24 maize and 37 Arabidopsis RALFs. The generated tree was combined with expression data from five tissues including pollen (Figure 1A). Predicted signal peptide and pro-peptide were removed and only mature protein sequences were used for comparison. 17 out of 24 maize RALFs belong to Clade IA and Clade IB, while the remaining 7 maize RALFs belong to Clade III. While Clade IA contains mainly vegetatively expressed *RALFs* from both species, Clade IB contains pollen-expressed *AtRALF4/19* and *ZmRALF2/3* representing their closest maize homologs that are also strongly expressed in pollen. Another maize homolog, *ZmRALF7*, only shows moderate expression in pollen (Figure 1A). Clade II contains only Arabidopsis *RALFs*, of which a few are expressed in pollen, while Clade III contains members from both plant species of which several genes show expression in pollen and *ZmRALF1/5* belong to this clade. The structural difference between the mature RALF proteins is visualized in Supplemental Figure S2. Clade IA and Clade IB RALFs contain typical RALF structures (Campbell and Turner, 2017) including YISY and YY interaction domains and four conserved cysteines. Clade IA RALFs show strong N-terminal variation, while Clade IB RALFs are less conserved at their C-termini. The YISY box, but not the YY motif is still recognizable in Clade II RALFs. Both motifs disappeared in Clade III RALFs together with the first two conserved cysteines.

**Figure 1.**
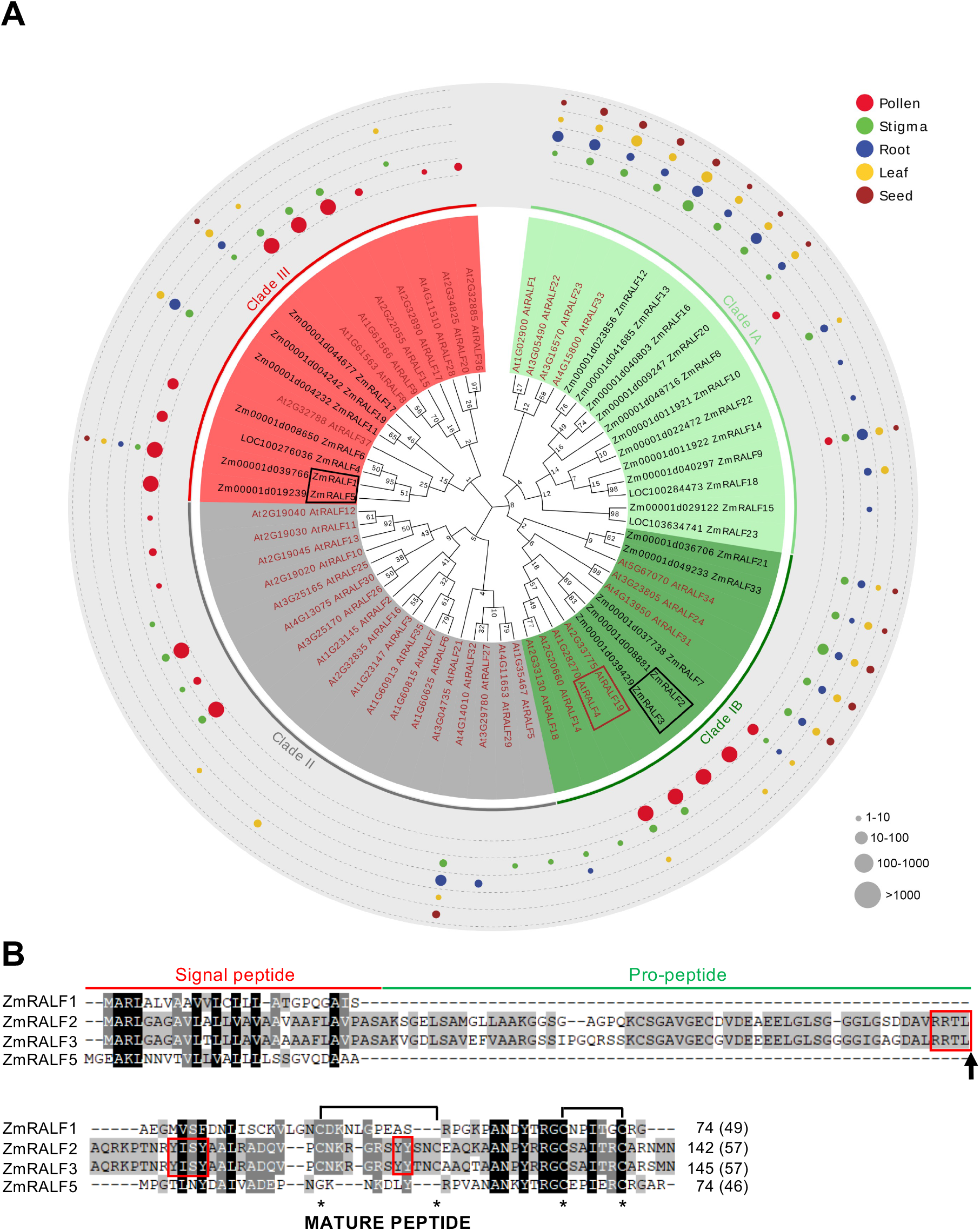
Phylogenetic and expression analysis of maize and Arabidopsis RALF protein families. **(A)** Phylogram using predicted mature maize (24 genes) and Arabidopsis (37 genes) peptide sequences. Arabidopsis AtRALFs are indicated in red and maize ZmRALFs in black. RALFs being investigated in more detail in this study are highlighted using rectangles. Clades IA and IB indicate RALFs with conserved RXXL/RXLX S1P protease cleavage sites and YISY as well as YY domains, which are lacking in Clade II and Clade III RALFs. Colored dots show expression pattern of *RALF* genes in the five selected tissues indicated. Dot size correlates to expression level determined by TPM (transcript per million) values as indicated. See Supplemental Figure S1 for details about expression values and Supplemental Figure S2 for amino acid conservation in Clade I-III RALFs. **(B)** Amino acid alignment of ZmRALF1/2/3/5 proteins. RRTL S1P protease cleavage site (arrow) and YISY as well as YY interaction domains of Clade I RALFs are boxed in red. Cysteine residues are marked with asterisks. Intramolecular disulfide bridges are indicated. Length of predicted mature peptides is indicated in brackets behind full length amino acid numbers.

Protein sequence alignment of the four most strongly expressed RALFs from maize pollen shows that ZmRALF2/3 contain typical RALF structures (Figure 1B; Supplemental Figure S2). These include a RXXL/RXLX (X represent any amino acid) motif for cleavage by SITE-1 PROTEASE (S1P) to remove the pro-peptide and to generate the mature peptide (Matos et al., 2008; Stegmann et al., 2017), four conserved cysteine residues to fold a proper structure, a YISY motif as part of an N-terminal α-helix for interaction with LLG co-receptors as well as the YY double tyrosine amino acid motif and partly conserved C-terminal region for likely interaction with CrRLK1L receptor kinases (Xiao et al., 2019). Based on structural studies using their close homolog AtRALF4 (Moussu et al., 2020), it is further likely that ZmRALF2/3 are capable to interact with LRX/PEX cell wall proteins. In contrast, the highest expressed *RALF* gene in maize pollen encoding ZmRALF1, as well as ZmRALF5 lack those motifs, suggesting that these RALFs may have different functions during maize pollen germination and growth. Such genes were previously identified as *RALF-like* genes (Campbell and Turner, 2017), although they still fit the model of the RALF family (Silverstein et al., 2007). Thus, we also name them as *ZmRALFs* throughout the manuscript.

### Downregulation of Pollen-Specific *RALFs* Leads to Pollen Tube Burst and Male Gamete Transmission Reduction

To investigate the function of pollen-specific *RALFs* in maize, a *RNAi-ZmRALFs* gene silencing construct was generated based on *ZmRALF1/2/3-specific* sequences (see Materials for details) and transformed into maize. As shown in Supplemental Figure S3, *ZmRALF1/2/3* were downregulated to different levels in independent *RNAi-ZmRALFs* mutant lines. Line1, Line5 and Line7 showed a reduction of transcript levels to about 50% compared with that of corresponding wild type (WT) plants and were thus selected for further phenotypic analyses. Pollen germination and growth were compared *in vitro* between *RNAi-ZmRALFs* and regenerated plants lacking the transgene (used as WT control). On average, pollen germination ratio was very similar in *RNAi-ZmRALFs* lines (65.71%) and WT plants (65.68%). Following pollen tube growth, pollen tubes from RNAi-*ZmRALFs* mutant lines started to burst, while burst of WT pollen tubes was initially rare (Figure 2A). A detailed analysis showed that burst of pollen tube occurred exclusively at the sub-apical region but never at the very tip (Figure 2B, C). Some pollen tubes even continued their growth despite burst and loss of significant cytoplasmic contents (Figure 2C). At 80 minutes after germination *in vitro*, more than 30% of pollen tubes from *RNAi-ZmRALFs* lines were burst, while only 7.2% WT pollen tubes burst at identical conditions.

**Figure 2.**
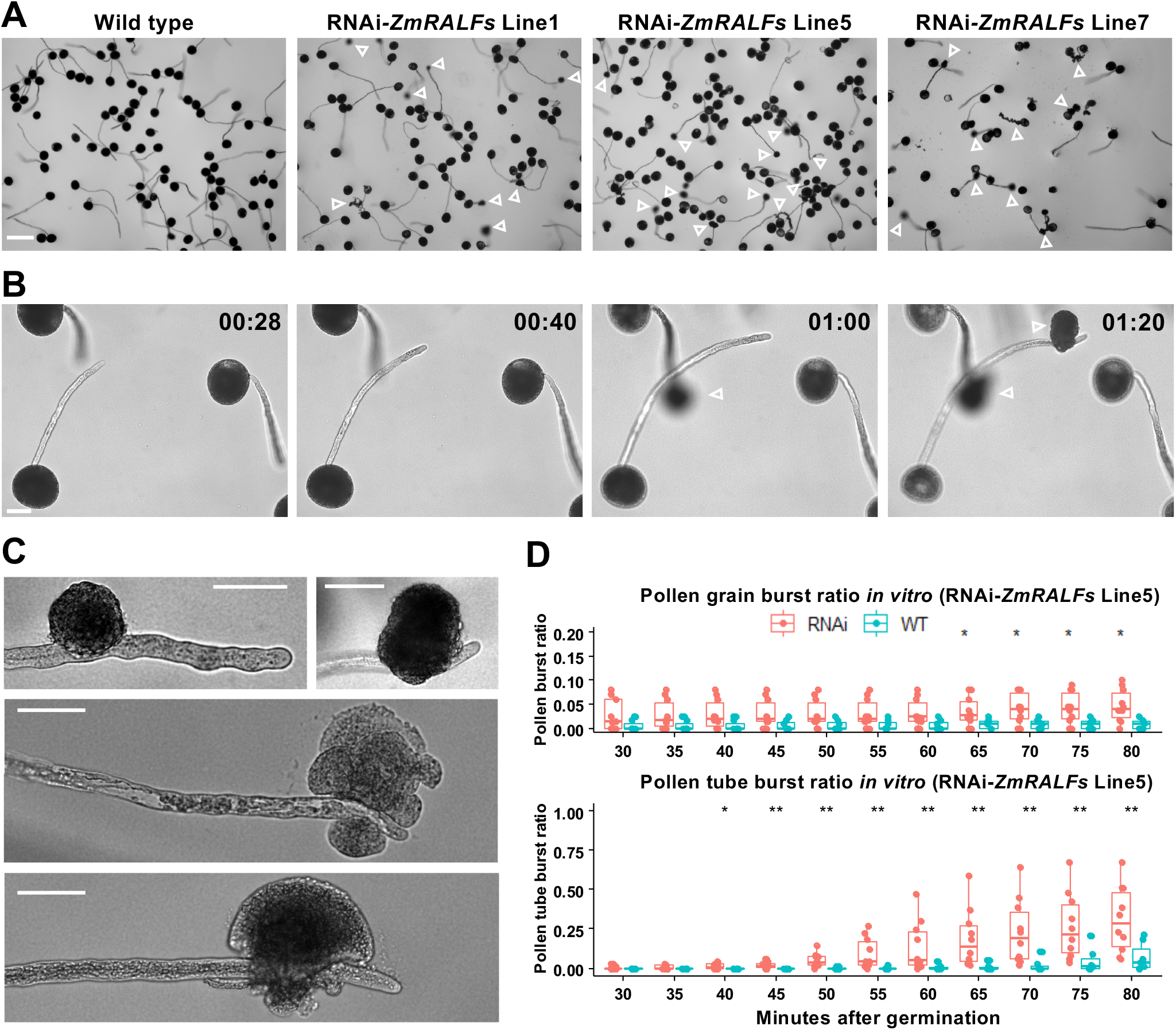
Pollen tubes of *RNAi-ZmRALFs* mutant maize lines burst *in vitro*. **(A)** Germination and growth of pollen from wild type and three RNAi-*ZmRALFs* lines after 1 hour. Open white arrowheads point towards burst pollen tubes. **(B)** Time-lapse imaging of pollen from RNAi-*ZmRALFs* mutant Line5 at indicated time points. One pollen tube burst at 60 and the second one after 80 minutes. Open white arrowheads point towards burst pollen tubes. **(C)** Examples of burst pollen tubes. Note that all pollen tubes burst at the shank but never at the apex of the tube. **(D)** Burst ratio of pollen from RNAi-*ZmRALFs* mutant Line5 was determined over time. Un-germinated pollen grains (top) and germinated tubes (bottom) were recorded. Scale bars are 200 μm in A and 50 μm in B and C. Significance: * p<0.05, ** p<0.01.

To further explore whether reduction of *ZmRALFs* expression levels in mutant pollen has also an effect *in vivo*, we determined male and female transmission efficiency of RNAi-*ZmRALFs* mutant lines (Table 1). *RNAi-ZmRALFs* mutant lines were either crossed as maternal or paternal parent with wild type plants (inbred line B73). Because transgenic lines often contain multiple integrations, we calculated an expected segregation ratio of ≥1:1 when RNAi-*ZmRALFs* lines were crossed with WT plants and ≥3:1 in self-crosses of *RNAi-ZmRALFs* mutant lines. As shown in Table 1, when Line1 and Line5 were used as pollen donor, transmission efficiencies of 0.72 and 0.24, respectively, were obtained. This is significantly lower than the expected number of ≥1. Similarly, transmission efficiencies of 0.69 and 0.44, respectively, were observed during self-crossings, which is much lower than the expected number of ≥3. In contrast, when we used the same lines as female parent, transmission efficiencies of 0.94 and 1.0, respectively, were obtained, which are very close to the expected number of ≥1. These genetic studies demonstrate that reduction of *ZmRALFs* levels in pollen affects male, but not female gamete transmission.

**Table 1.** Transmission efficiency of *RNAi-ZmRALFs* mutant maize lines.

### Only Clade IB ZmRALFs Locate to the Cell Wall *in vitro*

Previous reports showed that GFP-tagged *NtRALF* was detected first in the ER and later in the cell wall of epidermal cells (Escobar et al., 2003), which is similar to pollen-expressed AtRALF4/19 that were mainly observed in the apoplast (Ge et al., 2017) and cell wall (Mecchia et al., 2017). Thus, to explore subcellular localization of Clade IB and Clade III ZmRALFs, transgenic Arabidopsis plants expressing the four *ZmRALFs* used in this study were generated. Around 2 kbp genomic sequence upstream of the ORF of *AtRALF4* was taken as promoter and eGFP was cloned C-terminally to *ZmRALF*-ORFs. Arabidopsis pollen tubes from at least three independent transgenic lines were observed by using propidium iodide (PI) for counter staining of the cell wall. As shown in Figure 3 A-G, ZmRALF2/3-eGFP expressing pollen tubes display a slight overexpression phenotype indicated by their bumpy growth behavior. Their eGFP fusion proteins are visible in the ER, in small cytoplasmic vesicles and finally accumulate at the pollen tube tip and the cell wall showing co-localization with PI. Thick cell wall regions accumulate larger quantities of ZmRALF2/3-eGFP during transient growth retardation. In contrast, ZmRALF1/5-eGFP could be observed in the ER and larger cytoplasmic vesicles or granules, but never in the very tip of the pollen tube or cell wall, indicating that they are not secreted to the apoplast *in vitro*.

**Figure 3.**
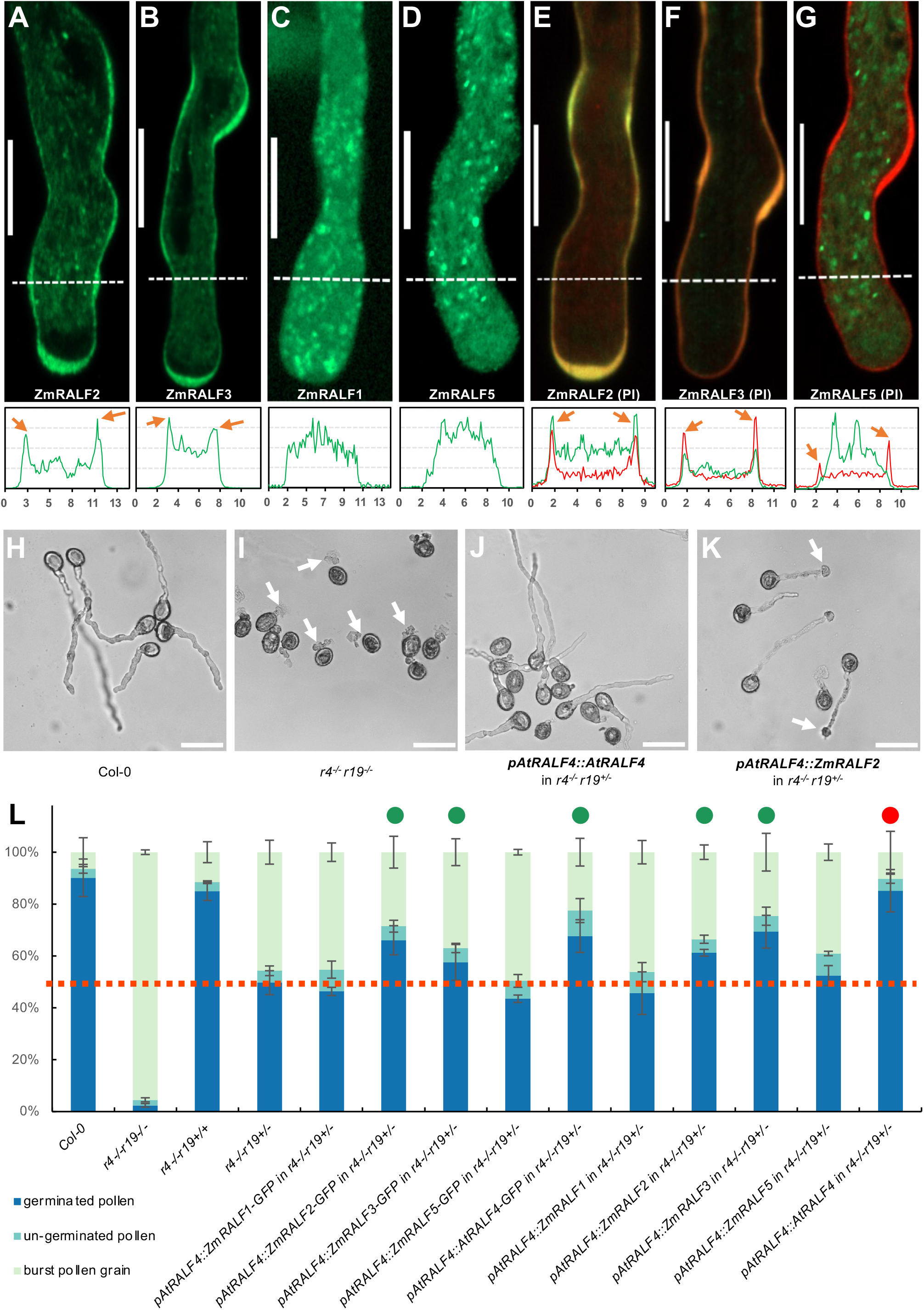
Localization and complementation studies of maize Clade IB and Clade III RALFs in Arabidopsis pollen tubes. **(A-G)** Around 2 kb upstream of the *AtRALF4* gene was used as promoter to express maize RALF proteins fused C-terminally to eGFP in Arabidopsis pollen tubes. Top row shows images of indicated ZmRALFs with or without propidium iodide (PI) counter staining. Bottom row shows correlated relative green and/or red fluorescence intensity each 10 μm from the pollen tube apex (illustrated by white dashed lines). The x-axis values represent the distance in μm from the first dash at the left along the dashed line and the y-axis shows relative fluorescence intensity. Arrowheads indicate fluorescence signal enrichment in the cell wall. Scale bars are 10 μm. **(H)** *In vitro* growth of WT pollen tubes. **(I)** Burst of *AtRALF4/19* double homozygous mutant (*r4*^-/-^ *r19*^-/-^) pollen tubes immediately after germination. Arrows point towards burst pollen tubes. **(J)** *AtRALF4* can fully restore *in vitro* germination and growth of the *r4*^-/-^ *r19*^+/-^ mutant pollen. Scale bars are 50 μm in H, I, J, K. **(K)** Pollen tubes from a partially complemented *ZmRALF2* transgenic plant showed partial rescue of germination and a late bursting phenotype in the *r4*^-/-^ *r19*^+/-^ mutant background. Arrows points towards burst pollen tubes. **(L)** Quantification of *in vitro* pollen germination and burst ratio in transgenic lines indicated. Red dotted line indicates germination ratio of 50%. Green dots represent transgenic lines that partially complemented the mutant *r4*^-/-^ *r19*^+/-^ phenotype. The red dot indicates *AtRALF4* that fully rescued the pollen grain burst phenotype of the *AtRALF4/19* double mutant.

### Clade IB *ZmRALFs* are Partially Capable to Complement Arabidopsis *ralf4/19* Mutants

Although ZmRALF2/3 belong to the same clade as AtRALF4/19, it was unclear whether they are able to functionally complement each other *in planta*. Therefore, above RALFs from maize and AtRALF4 from Arabidopsis as a control were expressed with and without eGFP fusions in the *ralf4/19* mutant background. As shown in Figure 3H, WT Col-0 pollen germinated well *in vitro* on solid germination medium with a germination ratio of around 90%, only a small portion of pollen grains burst (Figure 3L). As reported before (Ge et al., 2017), almost all pollen grains of the double homozygous *ralf4/19* mutant burst immediately after germination (Figure 3I, L). Due to gene redundancy, pollen germination of single mutants behaved similar to the WT (Figure 3L), and around half of the pollen grains from the heterozygous *ralf4-/-19+/-* mutant burst, while the other half germinated normally (Figure 3L). Full complementation was observed using AtRALF4 (Figure 3J, L), while addition of eGFP reduced the complementation ratio (Figure 3L). ZmRALF2/3 can partially restore the germination burst phenotype, but pollen tube burst was observed during further growth (Figure 3K). The eGFP-tag had no influence on ZmRALF2/3 complementation of immediate pollen tube burst (Figure 3L). In contrast, Clade III ZmRALF1/5 were not capable to rescue the pollen germination burst phenotype (Figure 3L). In conclusion, although ZmRALF2/3 and AtRALF4/19 are the closest homologs (Figure 1A), ZmRALF2/3 can only partially complement the *ralf4/19* mutant. Together with above findings showing that *ZmRALF* mutants show a pollen tube growth but not a pollen germination defect in maize, these data indicate functional diversification of pollen-specific Clade IB RALF genes in different plant species.

### Pollen Tube Integrity is Regulated by *ZmRALF2/3*

Based on phylogenetic analyses, subcellular localization pattern and complementation results, we assumed that Clade IB and Clade III *ZmRALFs* possesses distinct functions in pollen germination and pollen tube growth. To investigate the function of the two clades independently, CRISPR-Cas9 editing mutants were generated. After maize transformation, crossing and genome editing pattern analysis, Cas9-free homozygous mutant lines were used for *in vitro* pollen germination tests. As shown in Figure 4A and B, two independent transgenic events from each transformation with different genome editing patterns were selected for further phenotypic analysis. In *ZmRALF1/5-Cas9* line1, a 1 bp deletion in *ZmRALF1* caused a pre-stop codon generating a truncated peptide of 34 aa (instead of 74 aa still including a signal peptide of 25 aa). A 2 bp insertion in *ZmRALF5* caused a frame shift mutation generating a non-sense peptide sequence of 146 aa instead of 74 aa (including a signal peptide of 28 aa). In *ZmRALF1/5-Cas9* line2, a 2 bp deletion in *ZmRALF1* caused a pre-stop codon generating a truncated peptide of 34 aa, and a 1 bp insertion in *ZmRALF5* generated a shortened peptide of 69 aa. In *ZmRALF2/3-Cas9* line1, a 1 bp insertion in *ZmRALF2* caused a pre-stop codon generating a truncated peptide of 52 aa (instead of 142 aa) excluding the predicted mature peptide sequence and a 1 bp insertion in *ZmRALF3* causing a frame shift mutation generated a 203 aa non-sense peptide lacking the predicted mature peptide sequence. In *ZmRALF2/3*-Cas9 line2, a 1 bp insertion in *ZmRALF2* caused a frame shift mutation generated a 183 aa non-sense peptide lacking the predicted mature peptide sequence. A 21 bp deletion in *ZmRALF3* caused the lack of the 8 aa SSIPGQRS sequence in the pro-peptide that was replaced by G. As consequence, the mature peptide sequence was kept leading to a single *ZmRALF2* mutation.

**Figure 4.**
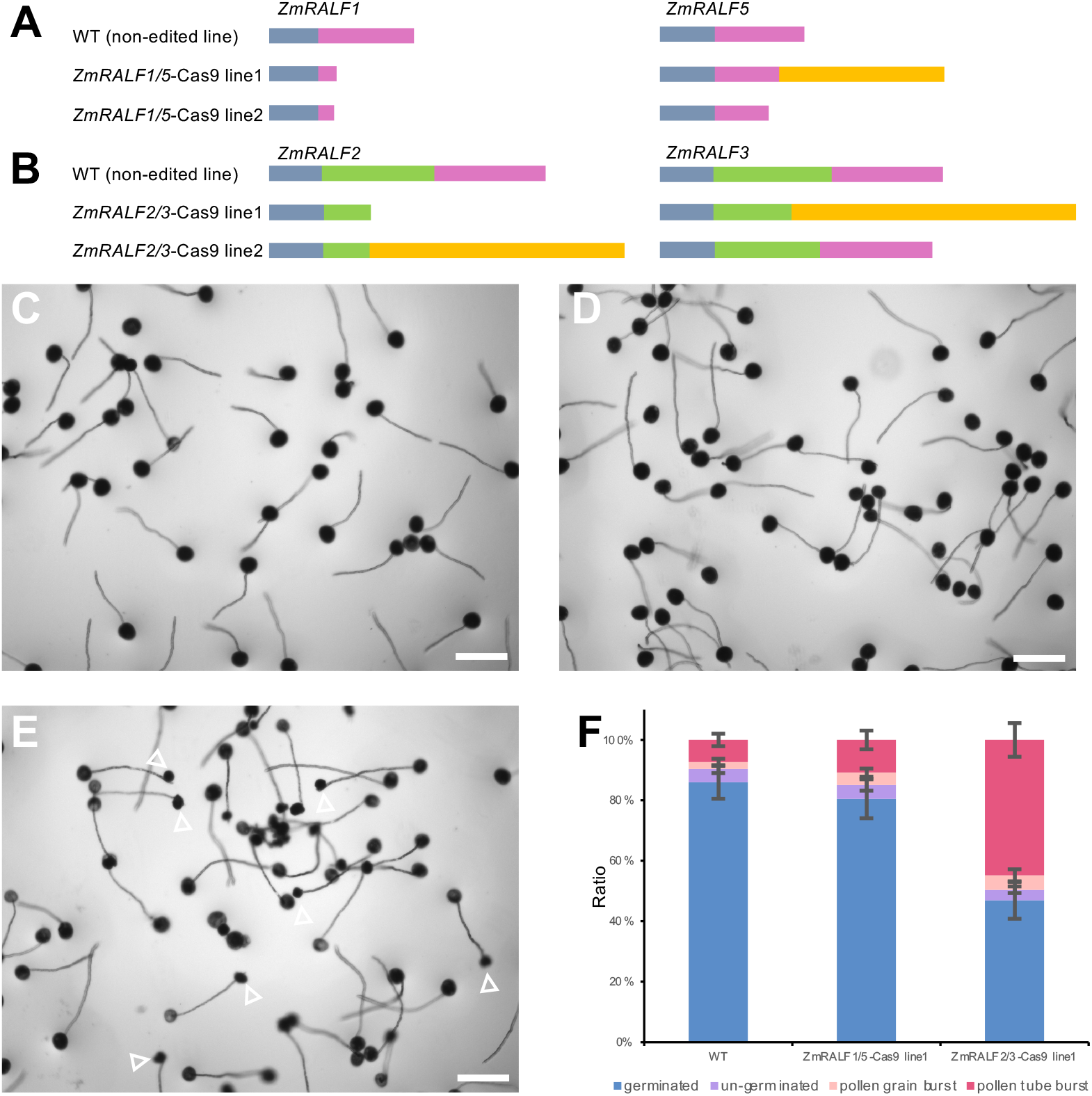
Editing pattern and phenotypic analysis of *ZmRALF1/5-Cas9* and *ZmRALF2/3-Cas9* mutant lines. **(A)** Genome editing pattern of *ZmRALF1* and *ZmRALF5* in mutant lines indicated. Blue-grey boxes show N-terminal signal peptides and pink boxes predicted mature peptides. Orange box indicates nonsense sequence. **(B)** Genome editing pattern of *ZmRALF2* and *ZmRALF3* in mutant lines indicated. Color code of boxes as in (A). Green box additionally shows the cleaved pro-peptide sequence (see also Fig. 1B). **(C-E)** *In vitro* pollen germination of WT (non-edited transgenic line) in (C), *ZmRALF1/5-Cas9* line1 (D), and *ZmRALF2/3-Cas9* line1 (E), respectively. Pollen grains were germinated on PGM for 60min. Arrowheads in (E) point towards burst pollen tubes. Scale bars are 200 μm. **(F)** Quantification of pollen burst ratios in WT (non-edited transgenic line), *ZmRALF1/5-Cas9* line1 and *ZmRALF2/3-Cas9* line1, respectively. n>200 in each plant. SD stands for difference from three independent pollen germination experiments.

During *in vitro* pollen germination tests, *ZmRALF1/5-Cas9* lines behaved similarly to WT plants, while pollen tube from *ZmRALF2/3*-Cas9 lines burst during growth (Figure 4C-F), which is very similar to *ZmRALFs*-RNAi lines reported above. This result clearly indicates that during *in vitro* pollen germination, *ZmRALF2/3* from Clade IB contribute to the regulation of pollen tube integrity maintenance while *ZmRALF1/5* from Clade III have no obvious impact suggesting that *ZmRALFs* from Clade IB and III have distinct functions during pollen tube growth.

### Pollen-Specific CrRLK1Ls ZmFERL4/7/9 Interact with Clade IB RALFs

In Arabidopsis it was shown that AtRALFs interact with *Catharanthus roseus* receptor-like kinases (CrRLK1Ls). The most prominent member of the family, FER, which is not expressed in pollen, interacts with root- and stigma-expressed Clade IA AtRALF1/23/33 (Haruta et al., 2014; Liu et al., 2021) as well as pollen- and ovule-expressed Clade IB AtRALF4/19/34 (Gao et al., 2022). Further CrRLK1L members including THESEUS1 and pollen-specific ANX1/2 and BUPS1/2 were reported as Clade IB AtRALF4/19/34 receptors (Ge et al., 2017; Mecchia et al., 2017; Gonneau et al., 2018). Notably, a recent report showed that also Clade II AtRALF6/7/16 and Clade III AtRALF36/37 act via FER, ANJ and HERK1 receptor kinases (Zhong et al., 2022). To explore the CrRLK1L family of maize, above proteins from Arabidopsis were used as a query to BLAST the maize genome. We identified 17 CrRLK1L genes in maize (Supplemental Figure S4), which is identical with the number of genes in Arabidopsis. According to its founding member, we named all maize CrRLK1Ls as FER-Like (FERL). *ZmFERL1/2/3* genes show a broad expression in vegetative tissues and likely encode the FER orthologs. Three *ZmFERLs* are strongly and specifically expressed in pollen. According to its sequence identity and expression pattern, ZmFERL4 appears as the single ortholog of Arabidopsis ANX1/2, while ZmFERL7/9 show highest homology to BUPS1/2. It should be noted that the kinase domain of ZmFERL7 is shortened compared with the homologs from Arabidopsis (Supplemental Figure S5) and it thus may not be functional. In summary, it seems that the CrRLK1L family is highly conserved in maize and Arabidopsis, and all Arabidopsis CrRLK1L members have close homologs in maize. However, MDS1-4 lack homologs are lacking in maize and ZmFERL16/17 miss in contrast to their closest homologs a kinase domain indicating variation and differences in CrRLK1L signaling in both species.

Phylogenetic relationship and expression of pollen-specific ZmFERLs is summarized in Figure 5A. To investigate whether N-terminal-tagged ectodomains of pollen-specific ZmFERL4/7/9 as well as non-pollen expressed ZmFERL1 are capable to interact with pollen-expressed Clade IB and Clade III ZmRALFs, pull-down assays were performed with C-terminal His-tagged mature ZmRALF1/2/3/5 peptides. As shown in Figure 5B-D, Clade IB ZmRALF2/3, but not Clade III ZmRALF1/5 interact with all ZmFERLs being tested. To further uncover the preference of Clade IB ZmRALFs binding towards different ZmFERLs, microscale thermophoresis (MST) assays were performed. While ZmFERL1 showed lowest binding affinity (Kd of 810 nM with ZmRALF2 and 1733 nM with ZmRALF3), ZmFERL9 showed strongest affinity (Kd of 104-107 nM with both ZmRALFs), while ZmFERL4/7 interacted slightly weaker with both ZmRALFs with Kds between 253-374 nM (Figure 5E, F). On average, the binding affinity between pollen-specific ZmFERLs was more than five times higher compared with that of the non-pollen expressed FER homolog ZmFERL1. Next, we tested whether ZmFERLs are also capable to interact with Clade IB RALFs from Arabidopsis and *vice versa*. As shown in Supplemental Figure S6A and B, AtRALF4 interacted with all four maize ZmFERLs being tested, while AtRALF19 showed only a strong interaction with ZmFERL1. Similarly, ZmRALF2 interacted stronger with ANX1/2 and BUPS1/2 compared with ZmRALF3, and Clade III ZmRALF1/5 did not interact with Arabidopsis CrRLK1Ls (Supplemental Figure S6C). A very weak interaction was observed between ZmRALF1/5 and BUPS2. These findings indicate binding preference between RALFs and CrRLK1Ls from maize and Arabidopsis.

**Figure 5.**
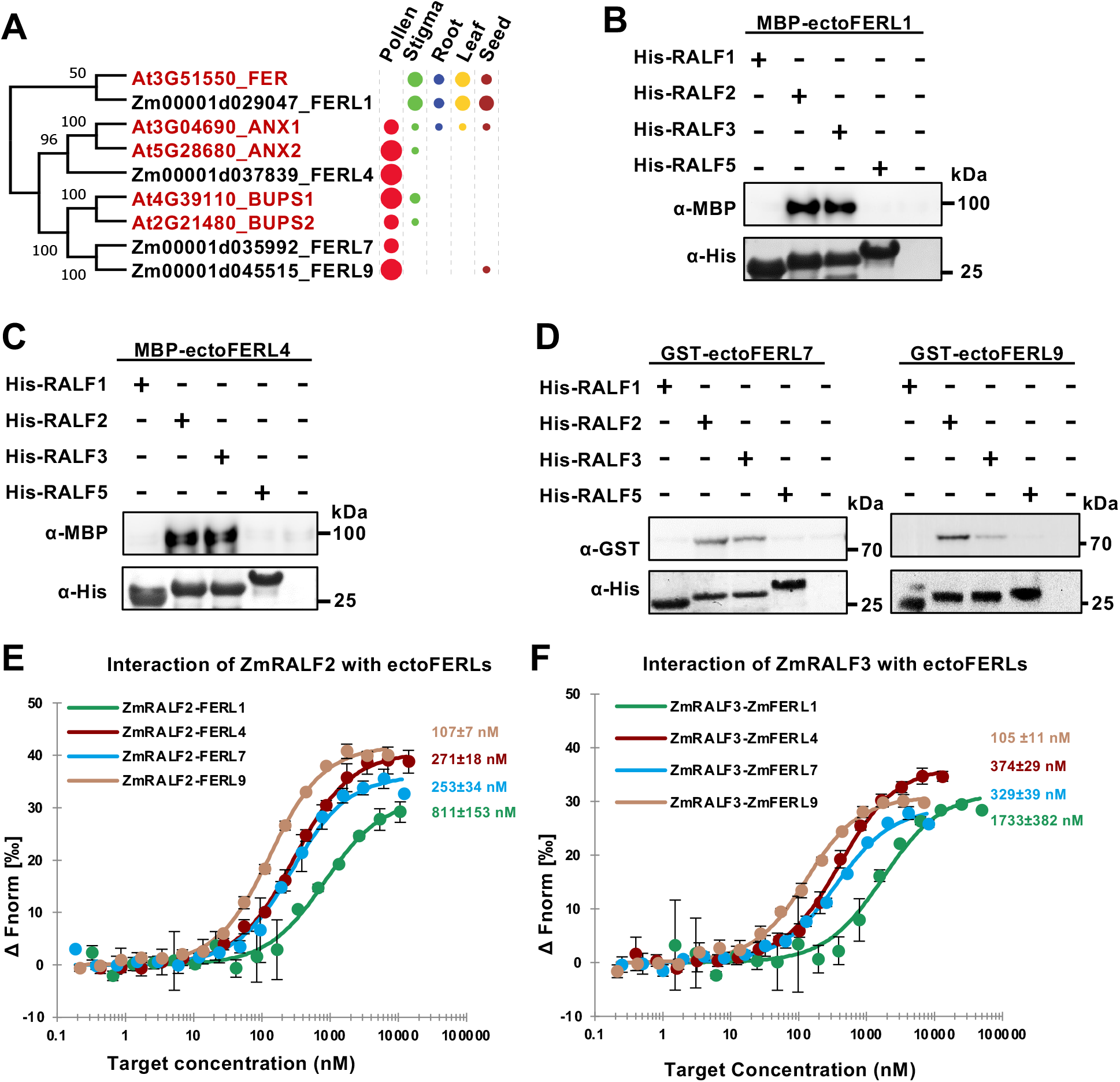
Clade IB, but not clade III pollen tube-expressed RALFs interact with CrRLK1L receptor kinases. **(A)** Phylogenetic analysis of ectodomains of selected Arabidopsis and maize FERONIA-like (FERL) receptor kinases. Expression pattern and relative transcript levels are indicated by dots (see Fig. 1A for explanations). Supplemental Figure S4 lists all maize and Arabidopsis CrRLK1Ls and shows detailed expression values. **(B-D)** Pull-down assays using MBP-tagged ectodomains of the FER homolog ZmFERL1 (B), the ANX1/2 homolog ZmFERL4 (C) as well as GST-tagged ectodomains of the BUPS1/2 homologs ZmFERL7/9 (D) each in combination with His-tagged ZmRALF1/2/3/5 as indicated. Only ZmRALF2/3 interact significantly with CrRLK1Ls. **(E-F)** Microscale thermophoresis (MST) binding affinity assays using ZmRALF2/3 in combination with ZmFERL1/4/7/9. Kd values are indicated. Notably, lowest binding affinity occurs with the FER homolog ZmFERL1, which is not expressed in pollen.

### Clade IB RALFs Interact with LORELEI-like GPI-Anchored Proteins

It was further shown in Arabidopsis that pollen expressed LORELEI-like GPI-anchored protein 2 (AtLLG2) and AtLLG3 act as co-receptors in RALF-CrRLK1L interaction complexes (Ge et al., 2019c). Structural analysis revealed that the N-terminal domain of mature Clade IB RALFs functions as a glue to generate a FER-RALF23-LLG2 receptor complex in Arabidopsis (Xiao et al., 2019). In maize, we identified four LORELEI-like GPI-anchored proteins named as ZmLLG1-4. *ZmLLG1/2* display a pollen-specific expression pattern (Figure 6A), which are similar to *LLG2/3* in Arabidopsis (Ge et al., 2019c). Notably, *ZmLLG3/4* expressed in vegetative tissues are slightly more similar to *LLG2/3* compared with *ZmLLG1/2*. Both Clade IB ZmRALF2/3 interacted with ZmLLG1/2, while interaction with Clade III ZmRALF1/5 was not observed (Figure 6B). Sequences alignment revealed conservation among LLGs from Arabidopsis and maize, respectively (Figure 6C). Especially the region covering eight cysteines is highly conserved including an Asn-Asp (ND) domain and two glycines between 7th and 8th cysteines that were reported to represent the core responsible for RALF binding in Arabidopsis (Xiao et al., 2019). We thus assume that all maize ZmLLGs are also capable to interact with ZmRALFs in a similar manner. Notably, ZmLLG1 lacks the C-terminal omega-side and GAS-domain for GPI-anchor addition indicating that it is likely not located at the surface of the plasma membrane. We thus suppose that ZmLLG2, which interacted stronger with Clade IB ZmRALFs compared with ZmLLG1, and whose gene shows also a stronger expression level, represents the major maize homolog of pollen-expressed Arabidopsis LLGs.

**Figure 6.**
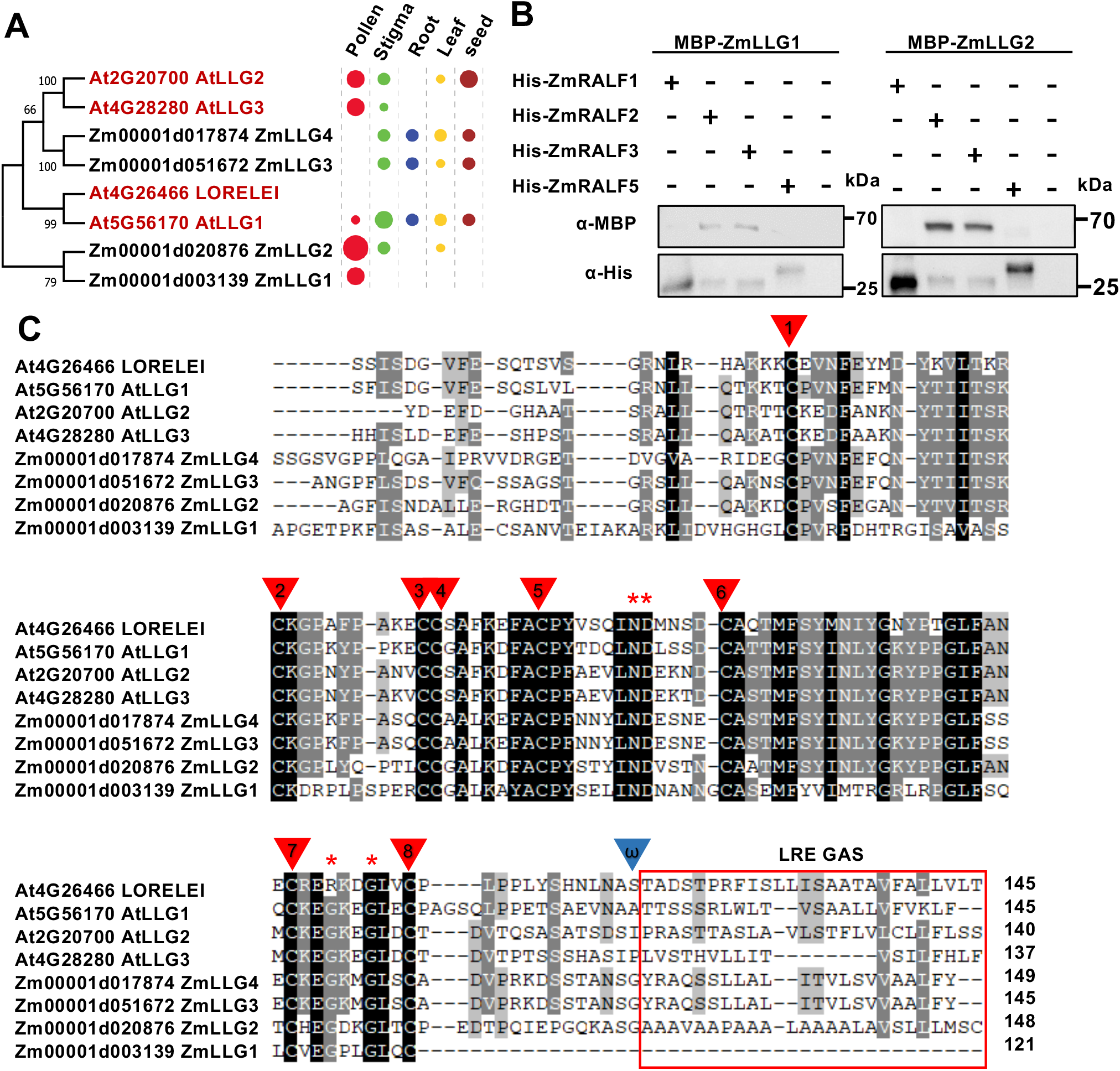
Clade IB maize RALFs are capable to interact with LORELEI-like-GPI-anchored (LLG) co-receptors. **(A)** Phylogenetic tree of LLG proteins from maize and Arabidopsis. Expression pattern and relative transcript levels are indicated by dots (see Fig. 1A for explanations). **(B)** Pull-down assay using MBP-tagged LLGs and His-tagged RALFs as indicated. **(C)** Alignment of maize and Arabidopsis LGGs (the numbers provide full-length sequences). Note that N-terminal signal peptide sequences were cut off for the alignment. Mature LGGs contain eight conserved cysteine residues (red arrowheads) and a conserved Asn-Asp (ND) domain and glycine (red asterisks). The region required for GPI-anchor addition (omega-side shown by a blue arrowhead and GAS domain boxed in red) are indicated at the C-terminus. This region is lacking in ZmLLG1.

### Clade IB ZmRALFs Interact with PEX Cell Wall Proteins and Regulate Their Phosphorylation Pattern

After elucidating that the ZmFERL4/7/9-ZmLLG2 receptor complex is likely involved in ZmRALF2/3-regulated pollen tube growth, we next aimed to identify downstream components of the signaling pathway. We expected that signaling components are either hypo- or hyperphosphorylated in response to receptor complex activation and compared the phosphorylation pattern of RNAi-*ZmRALFs* lines of pollen tubes grown for 45 min *in vitro* with that of WT plants. The phospho-proteome was determined by mass spectrometry. Surprisingly, the major protein group showing hyper-phosphorylation were ZmPEX leucine-rich repeat (LRR)-extensin-like proteins. Previous reports showed that the homologs of ZmPEXs cell wall proteins (named LRX in Arabidopsis) directly interact with RALFs and sustain pollen germination and tube growth via the CrRLK1Ls pathway (Mecchia et al., 2017; Moussu et al., 2020). Supplemental Figure S7 shows phosphoprotein intensity of ZmPEX2 as an example in two RNAi-*ZmRALFs* lines compared with WT controls. 26 possible serine/threonine phosphorylation sites were detected in the proline-rich extensin domain of ZmPEX2 and one in the LRR-domain (Figure 7A). Compared with ZmPEX2 in WT pollen tubes, phosphorylation sites were significant hyperphosphorylated in the RNAi-*ZmRALFs* lines and S288 in the LRR-domain was most strongly affected (Supplemental Figure S7B).

**Figure 7.**
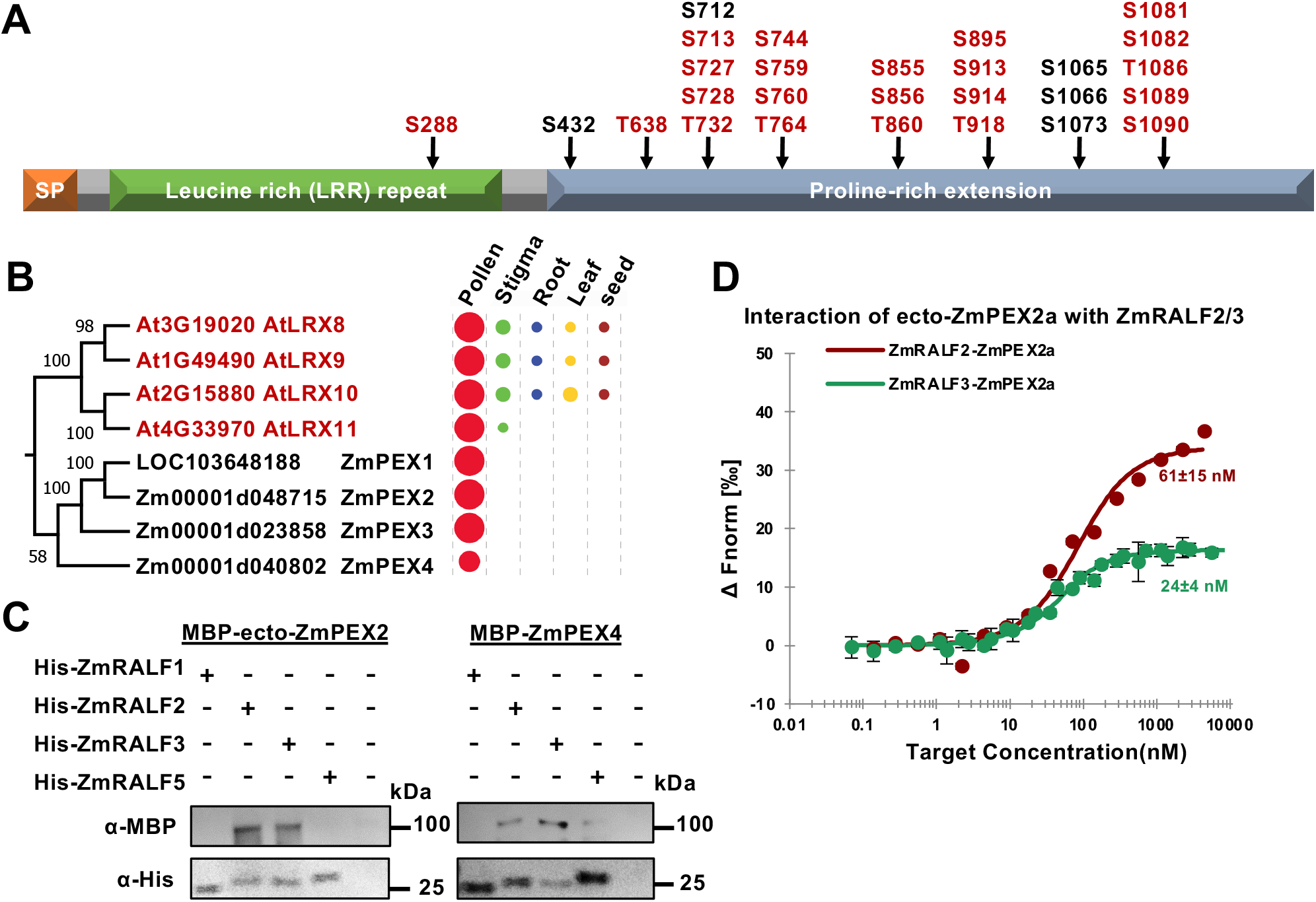
Cell wall-localized maize PEX proteins are hyper-phosphorylated in RNAi-*ZmRALFs* lines and physically interact via their LRR-domain with Clade IB RALFs. **(A)** ZmPEX (Pollen Extensin-Like) proteins were identified as major targets of pollen-specific RALF signaling in a screen for proteins with changed phosphorylation pattern in RNAi lines. Schematic diagram of the chimeric ZmPEX2 protein structure consisting of an N-terminal signal sequence (SP) for secretion, a globular leucine-rich repeat (LRR) and a proline-rich extensin-like domain. With the exception of ZmPEX4 lacking the extensin-like domain, all maize PEX and Arabidopsis LRX proteins shown below contain this protein structure. 27 putative serine (S) and threonine (T) phosphorylation sites are indicated. Hyperphosphorylated sites are colored in red. See Supplemental Figures S7 and S9 for details. **(B)** Phylogenetic analysis of pollen-specific chimeric LRR-extension-like proteins in maize and Arabidopsis. The signal peptide sequence and conserved proline-rich extensin domain were removed before alignment. Expression pattern and relative transcript levels are shown as explained in Figure 1A. The complete phylogenetic tree and detailed expression data including all LRR-extension-like proteins in maize and Arabidopsis as well as a protein alignment are shown in Supplemental Figures S8 and S9. **(C)** Pull-down experiment of MBP-tagged N-terminal LRR domain containing region of ZmPEX2 (ZmPEX2a) and full length ZmPEX4. **(D)** Microscale thermophoresis (MST) binding affinity assay showing that ZmRALF2/3 have strong binding affinity with ZmPEX2a: ZmRALF2, Kd = 60 nM; ZmRALF3, Kd = 24 nM.

Therefore, we next studied whether ZmRALF2/3 are capable to interact with ZmPEXs. Until now, it was only known that ZmPEXs localize towards the intine of mature maize pollen grains and the callosic sheath of pollen tubes (Rubinstein et al., 1995b; Rubinstein et al., 1995a). Their homologs in Arabidopsis were recently shown to interact with AtRALF4/19 and stabilize the pollen tube cell wall during germination by an unknown mechanism (Mecchia et al., 2017; Moussu et al., 2020). We used ZmPEX2 as a query and identified a total of 19 PEX/LRX genes in the maize genome and confirmed 11 LRX genes previously identified in Arabidopsis (Supplemental Figure S8). This family is thus slightly larger in maize compared with Arabidopsis. Similar to the four Arabidopsis homologs *AtLRX8-11*, the four *ZmPEX1-4* genes are specifically expressed in pollen (Figure 6A and 7B) with *ZmPEX3* belonging to one of the most strongly expressed genes in pollen. In addition to their N-terminal LRR domain, PEX/LRX proteins contain a conserved cysteine-rich motif to enhance homodimer formation (Moussu et al., 2020) and a proline-rich extension domain with multiple SPPPP repeats that extend to other cell wall components for anchoring (Herger et al., 2019). A sequence alignment of the eight PEX and LRX proteins generated in pollen of maize and Arabidopsis, respectively, is shown in Supplemental Figure S9. The N-terminal N-D domain, LRR domain and 11 cysteines are highly conserved, while the proline-rich C-terminal region shows variation between SPPPP repeats. ZmPEX1-3 are slightly larger compared with their four Arabidopsis homologs. ZmPEX4 lacks the extensin-like domain and thus is strictly speaking not a PEX/LRX protein despite the high homology of the N-terminal part of the protein.

Pull-down experiments were carried out with Clade IB and Clade III ZmRALFs with the N-terminal MBP-tagged LRR domain of ZmPEX2 and full-length ZmPEX4 (Figure 7C). Only ZmRALF2/3 interacted with both ZmPEX proteins. MST measurements showed that ZmRALF2 binds to the LRR-domain of ZmPEX2 with a kD of 60nM, while the binding affinity between ZmRALF3 and ZmPEX2 is even higher with a kD of 24nM. In summary, binding of Clade IB ZmRALFs to ZmPEX2 is on average more than five time higher compared to affinities detected with pollen-expressed cell surface receptors ZmFERL4/7/9 (Figure 5E, F) and thus explains their predominant subcellular localization in the cell wall (Figure 3A, B). Similar to the interaction with maize ZmPEX2 and ZmPEX4, Clade IB ZmRALFs were also capable to interact with pollen-expressed AtLRX9-11 from Arabidopsis (Supplemental Figure S10).

### Clade IB ZmRALFs Regulate Pectin Distribution and Wall Thickness in Growing Pollen Tubes

Proper biochemical composition and spatial distribution of cell wall material is critical for maintaining pollen tube growth as mutants defective in the synthesis of in cell wall components exhibit pollen tube growth phenotypes (Jiang et al., 2005; Nishikawa et al., 2005; Wang et al., 2011; Chebli et al., 2012). To investigate cell wall material distribution in the maize pollen tube, we first defined the different regions of growing pollen tubes. As shown in Figure 8A-B, the first ~3-4 μm of a growing pollen tube was defined as the “apex”, which is characterized by the clear zone extending from the outermost tip “pole”. The next ~5-10 μm is considered as the “sub-apex” region, and the “distal region” begins from ~10 μm from the tip.

**Figure 8.**
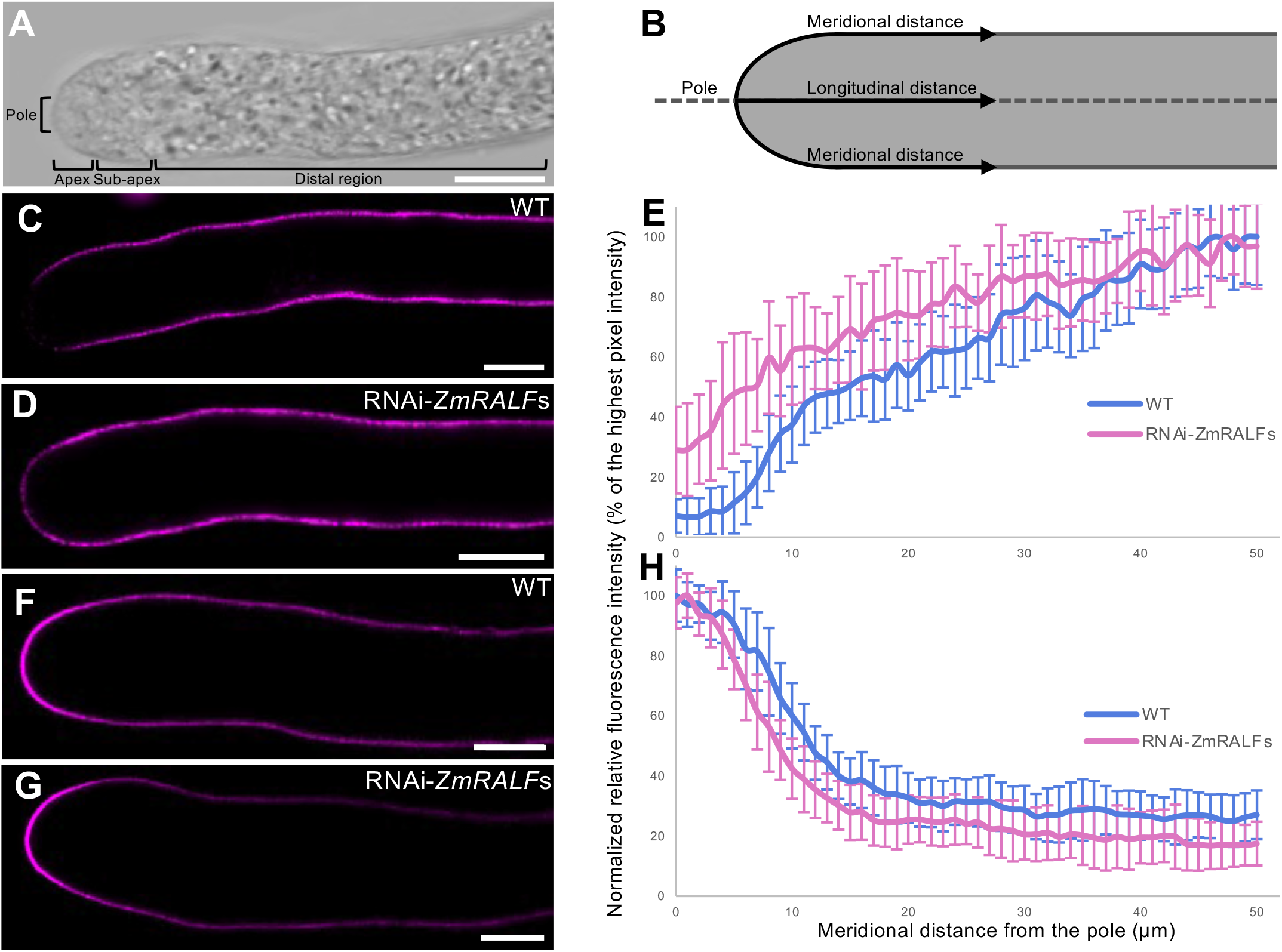
Pectin distribution in the pollen tube cell wall is altered in *RNAi-ZmRALFs* mutants. **(A)** Bright field image of a growing maize pollen tube showing various zones. The major growing zone is located at the pole and apex. The apex of the pollen tube is ~3-4 μm, while the sub-apical region is ~5-10 μm in length. ~10 μm from the pole begins the distal region of the pollen tube. **(B)** Schematic figure of a maize pollen tube. Fluorescent intensity in (C), (D), (F) and (G) was measured along meridional distances. **(C-D)** Representative images of JIM5 labelled WT and *RNAi-ZmRALFs* mutant pollen tubes, respectively. JIM5 monoclonal antibody labels de-esterified pectin/homogalacuronan. **(E)** Fluorescent intensity quantification of JIM5 labelled WT and *RNAi-ZmRALFs* mutant pollen tubes. n = 26 for both WT and *RNAi-ZmRALFs* pollen tubes, respectively. **(F-G)** Representative images of LM20 labelled WT and RNAi-*ZmRALFs* mutant pollen tubes, respectively. LM20 monoclonal antibody labels esterified pectin/homogalacuronan. **(H)** Fluorescent intensity quantification of LM20 labelled WT and RNAi-*ZmRALFs* mutant pollen tubes. For WT pollen tubes, n = 19 and for RNAi-*ZmRALFs* pollen tube, n = 21. Scale bars = 10 μm.

Pectins, which are also known as pectic polysaccharides, are the main and most important component of the pollen tube cell wall. Pectins are a heterogenous group of galacturonyl polymers containing contain galacturonic acid in the backbone that can be esterified or de-esterified at their carboxylic acid residues by methyl groups. Both forms can be labeled with JIM5 and LM20 antibodies that specifically recognize pectin molecules with low and high degrees of methyl esterification, respectively (Knox et al., 1990; Clausen et al., 2003; Verhertbruggen et al., 2009). As shown in Figure 8C and E, the amount of de-esterified pectin in WT maize pollen tubes is almost absent at the first 4-5 μm from the very tip of growing pollen tubes, then it slightly increases and reaches a plateau at around 40 μm along the meridional distance. In contrast, in RNAi-*ZmRALFs* mutant pollen tubes, de-esterified pectin can be detected from the very tip of the pollen tube and reaches similar levels to WT pollen from about 35 μm from the tip (Figure 8D and E). The amount of esterified pectin is similar in the apex of WT and RNAi-*ZmRALFs* mutant pollen tubes but decreases faster in the sub-apical region and remains lower in the distal region (Figure 8F-H).

Although different distribution patterns of de-esterified pectin and esterified pectin can be observed along mutant pollen tubes, it was unclear whether this distribution pattern may also affect the wall stiffness and/or thickness. Therefore, we optimized and finally increased the concentration of the fixative to facilitate its penetration into *in vitro* grown pollen tubes and to enhance the fixation rate. Significant differences in cell wall structure and thickness was detected in mutant pollen tubes compared with that of WT plants. As shown in Figure 9A and B, both esterified and de-esterified pectin labelled WT pollen tubes can be classified into two categories, (i) pollen tubes with normal and linear cell wall structure (Figure 9A and C) and (ii) pollen tube with split cell wall structure (Figure 9B and D), which caused cell wall thickness increasement (Figure 9E-H). Although splitting of cell walls can also be observed in WT pollen, the frequency increased from 40 to 80.8% in *RNAi-ZmRALFs* and even 92.9% in *ZmRALF2/3-* Cas9 mutant pollen tubes using the JIM5 antibody, while the observed frequency increased from 7.7 to 47.1% and 55.5%, respectively, by using the LM20 antibody (Figure 9A-D and Supplemental Figure S11). Moreover, the cell wall thickness in the sub-apex increased >30% in *RNAi-ZmRALFs* and *ZmRALF2/3-Cas9* mutant pollen from about 0.8-0.95 μm in WT pollen tubes to about 1.15-1.5 μm depending on the usage of the LM20 and JIM5 antibody, respectively (Figure 9I, J).

**Figure 9.**
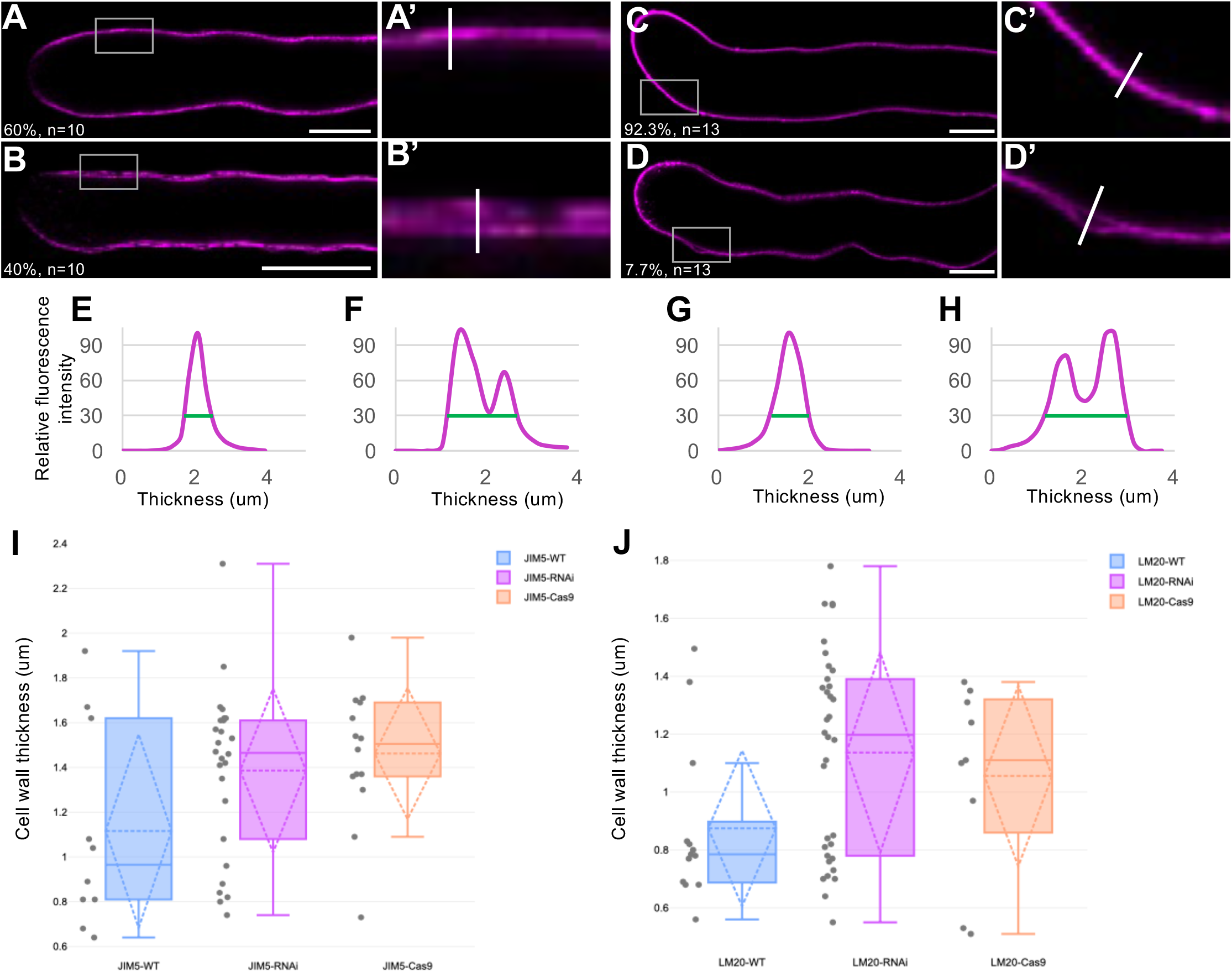
Cell wall organization pattern is altered in *ZmRALFs* mutants. **(A-B)** Representative images of JIM5 labelled WT pollen tubes. Cell walls either showed a thin homogenous line or were thicker and dilated showing partially two lines. The ratio of each category and the number of pollen tubes that were analyzed are indicated. (A’-B’) Enlargement of grey boxes in (A) and (B), respectively. Scale bars = 10 μm. See Suppl. Fig. S11A-D for *ZmRALF* mutant lines. **(C-D)** Representative images of LM20 labelled WT pollen tubes. The ratio of each category like in (A-B) and the number of pollen tubes that were analyzed are indicated. (C’ and D’) Enlargement of grey boxes in (C) and (D), respectively. Scale bars = 10 μm. See Suppl. Fig. S11E-H for *ZmRALF* mutant lines. **(E-H)** Quantification of relative fluorescence intensity of white lines shown in (A’-D’). Thickness of cell walls was measured with the criteria that relative fluorescence intensity was 30% (green lines). **(I-J)** Box plot analysis of pollen tube cell wall thickness WT and *ZmRALFs* mutant lines after labelling with JIM5 (I) and LM20 (J), respectively. The thickness was defined with the same criteria as in (E-H).

## DISCUSSION

So far, the RALF-LLG-CrRLK1Ls and RALF-LRX signaling pathways that play key roles during pollen tube germination, growth and sperm cell release have been studied exclusively in the model plant Arabidopsis and it is unclear to which extent obtained knowledge can be transferred to other plant species and how the pathways are connected. It is also unclear when and how these signaling pathways have been established during land plant evolution and when family members were further specified to take over specific roles in reproduction and pollen tube growth. *RALFs* and *CrRLK1Ls* were first detected in bryophytes. There exists, for example, a single *CrRLK1L* and three *RALF* genes in *Marchantia polymorpha* (Galindo-Trigo et al., 2016). A strong amplification of *RALF* genes was reported in angiosperms. While the basal angiosperm *Amborella trichopda* contains already nine genes, >30 genes were reported in various eudicot and monocot families (Campbell and Turner, 2017; Abarca et al., 2021). So far, research of *RALF* functions focused on eudicots and were especially conducted in Arabidopsis. In monocots, one study reported that in the grass sugarcane *SacRALF1* is involved in the regulation of tissue expansion (Mingossi et al., 2010). By comparing *RALFs* from the eudicot model Arabidopsis and the grass model maize, we distinguished three Clades. Our nomenclature differs from a previous phylogenetic study comparing 795 identified RALFs (Campbell and Turner, 2017). Four major clades we reported in that study with Clades I-III RALFs containing an RRXL protease cleavage site and the YISY motif for receptor interaction, while Clade IV contains all other RALFs. In the meantime, we have learned that the YISYxxLRRN domain mediates interaction with the LLG co-receptor forming a heterotrimeric complex with the CrRLK1L member FER (Xiao et al., 2019). Moreover, four tyrosine residues are involved in LRX-binding including the two Ys of the YISY motif and two additional ones in the YY motif in the middle of mature RALP peptides (Moussu et al., 2020). Notably, in the modified nomenclature suggested in this report, closely related Clade IA and IB contain all RALFs harboring above-described motifs, while these motifs are highly degenerated and lacking in Clade II and Clade III RALFs, respectively. Moreover, Clade IB RALFs that are more conserved after the cleavage site contain reproductive and pollen-specific RALFs (Ge et al., 2017; Mecchia et al., 2017; Gao et al., 2022) including RALFs of basal land plants like mosses, and thus appear to represent the most original clade. Clade IA, which is more variable after the cleavage site, contains mainly vegetatively and sporophytically expressed RALFs in the two analyzed species includes well described RALFs like AtRALF1, which may have acquired specific sub-functions during land plant evolution like regulation of root and root hair development (Haruta et al., 2014; Zhu et al., 2020) and AtRALF23/33 playing roles in papilla hair cell functions (Liu et al., 2021). Arabidopsis Clade II RALFs that don’t exist in maize and which still contain a degenerated YISY-motif, and even some members of Clade III, were recently shown to interact with CrRLK1L receptor kinases *in vitro* (Zhong et al., 2022). However, binding affinities were about three times lower compared to those measured for ZmRALF2/3 in this report. In contrast to above findings maize Clade III ZmRALF1/5 do not interact with CrRLK1Ls and likely acquired novel, unknown functions during evolution and the pollen tube journey through the specified transmitting tract tissue of the grasses (Zhou et al., 2017). It will now be important to study the role of Clade III ZmRALFs, but also to elucidate whether Clade II and III RALFs are still capable to interact with LLG co-receptors and LRX/PEX cell wall proteins.

Quantitative biochemical assays revealed that Clade IB AtRALF4 from Arabidopsis pollen binds LLGs and LRX cell-wall modules with drastically different binding affinities, and with distinct and mutually exclusive binding modes (Moussu et al., 2020). Binding of AtRALF4 with AtLRX8 could be very strong with a Kd of 3.5 nm, while strongest interaction with ANXs/BUPSs was reported at 310 nM (Ge et al., 2017). This drastic difference in binding affinity could be partially explained by optimized AtRALF4 folding in the first report, but points towards the same finding in the present study that ZmRALF2/3 bind on average about five times stronger to ZmPEXs compared to pollen-specific ZmFERLs, and even much weaker to ZmFERL1 which is not expressed in pollen. In conclusion, taking also into consideration that binding of LRX/PEX cell wall proteins and LGG-CrRLK1Ls receptor complexes to the RALF motif YISY motif is mutually exclusive, we propose the following model shown in Figure 10: at low RALF concentrations in the cell wall, RALFs bind almost exclusively to LRX/PEX cell proteins and the CrRLK1L-LLG signaling pathway is inactive. Similarly, if the cell wall is thick and solid while containing many LRX/PEX proteins, the signaling pathway is also silent. However, in the presence of high RALF levels in the pollen apex that contain thin cell walls, but also in thin cell wall of other tissues, the lower amounts of LRX/PEX proteins are saturated and surplus RALFs are capable to activate the CrRLK1L-LLG signaling pathway leading for example, to reduction of pollen tube growth speed or at high concentration to complete growth arrest allowing cells to secret sufficient cell wall material components and providing time to regulate the stiffness of the wall. If RALF levels are artificially reduced, the CrRLK1L-LLG signaling pathway remains silent even in thin cell walls and fine tune control of cell expansion, growth speed and wall stiffness that depends on cell wall integrity sensing leads to growth defects and in the case of pollen tubes to burst in the sub-apical region. Please note that this simple model did not consider further aspects, as (i) the described RALF interactions are also pH-dependent (Moussu et al., 2020) and (ii) its induced signaling modifies the redox status of the cell wall by RALF-induced generation of reactive oxygen species (ROS), which stimulates pollen hydration and tube growth while reducing the pollen burst rate (Feng et al., 2019; Liu et al., 2021).

**Figure 10.**
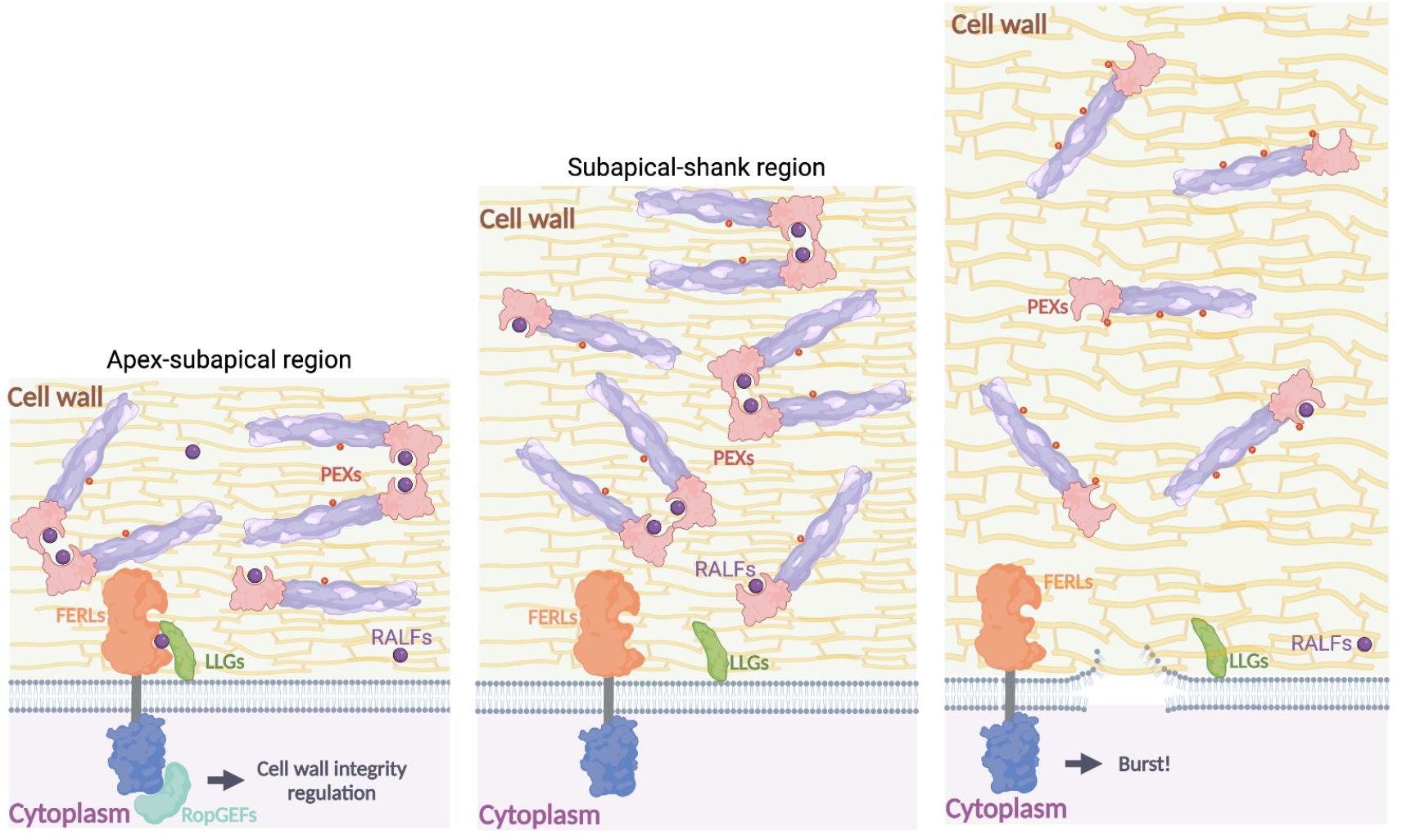
Dosage-dependent RALF signaling model showing the role of Clade IB RALFs as sensors of cell wall integrity and thickness during pollen tube growth. Left: during secretion of cell wall components at the growing pollen tube apex, the cell wall is thin and RALFs interact with PEX proteins and activate the FERL-LLG signaling pathway leading to further secretion of cell wall components, moderate phosphorylation of PEXs and wall stiffening. Middle: in sufficiently thick cell walls RALFs are predominately located at the cell wall due to higher affinity to PEX proteins that form dimers. The FERL-LGG signaling pathway becomes inactive. Right: the FERL-LGG signaling pathway is inactive in lines with reduced RALF levels. As consequences the cell wall is instable and dilated, and less PEXs form dimers. Hyper-phosphorylation of PEXs partially compensates cell wall instability but cannot prevent ultimate pollen tubes burst.

With the exception of above-described regulation of ROS levels and recently reported Ca^2+^ signaling (Gao et al., 2023), little is known about RALF-induced downstream signaling processes associated with pollen/pollen tube functions. In roots it has been shown that AtRALF1-FER interaction causes phosphorylation and thus inhibition of plasma membrane H^+^-ATPase2 explaining extracellular alkalinization (Haruta et al., 2014). AtRALF1 also enhances the interaction of FER with the receptor-like cytoplasmic kinase RIPK, which mutually phosphorylate each other to regulate cell growth in roots (Du et al., 2016). AtRALF1 treatment promotes direct phosphorylation of ERBB3 binding protein 1 (EBP1) by FER leading to its accumulation in the nucleus and inhibition of RALF peptide responses (Li et al., 2018). It was further shown that AtRALF1 promotes FER-mediated phosphorylation of the elongation factor eIF4E1. Phosphorylated eIF4E1 regulates the synthesis of root hair proteins including ROOT HAIR DEFECTIVE 6-LIKE 4 (RSL4), which is required for root hair growth (Zhu et al., 2020). Altogether these finding indicate that phosphorylation plays a key role in RALF-induced downstream signaling processes. Phospho-proteome measurements usually depend on RALF application and have to our knowledge neither been performed with pollen tubes nor with knockouts or RNAi lines. The finding that ZmPEXs are major targets of RALF signaling in maize and are hyperphosphorylated in *RNAi-ZmRALFs* lines was unexpected. It is unclear (i) whether RALFs interact with LRX/PEX proteins already in the secretory pathway or outside the plasma membrane in the cell wall and (ii) when LRXs/PEXs become phosphorylated. Notably, the phosphorylation site in the LRR-domain of ZmPEXs that was most often hyperphosphorylated may affect dimerization of ZmPEXs. This and perhaps other sites mighty be protected by RALFs bound already in the secretory pathway and thus prevent phosphorylation. Co-expression studies of labelled RALFs and LRX/PEX proteins will now be helpful to investigate whether interaction occurs already during the secretory pathway. It is also possible that an increase of phosphorylated ZmPEXs might counteract reduced RALF signaling and is used to add cross-bridges to pectic acids to partly stabilize the cell wall in *ralf* mutants. The generation of phospho-mimic mutants and structural studies would help to elucidate the role of LRX/PEX phosphorylation in future studies.

Together with callose pectins are the main components of the pollen tube cell wall, and their distribution pattern is of vital importance for maintaining proper pollen tube growth. In growing pollen tubes, high methyl-esterified (soft) pectins are deposited at the tip of pollen tube via exocytosis, and de-esterification takes place in the sub-apical region by the activity of pectin methylesterases (PMEs). The activity of PMEs in the very apical region is prevented by pectin methylesterase inhibitors (PMEIs) that are secreted within the same secretory vesicles (Scholz et al., 2020). By using immunolabelling of pollen tubes, we discovered that the spatial distribution of pectins is significantly altered under low RALFs level, and cell wall morphology is compromised ultimately leading to burst of pollen tubes. It remained unclear whether reduced RALF levels lead, for example, to faster endocytosis of PMEIs as indicated by higher levels of de-esterified pectins in the apex, to reduced ROS levels required to increase the stiffness of the wall in the sub-apical region or to reduced dimerization of ZmPEXs that likely contribute to stabilization of the cell wall. It is also unclear why pollen germination is not affected in the different ZmRALF2/3 mutants in maize and whether reduced RALF levels affect the Ca^2+^-gradient in the pollen tube apex that was reported to be dependent of RALF signaling (Gao et al., 2023). We assume that the latter is unlikely, otherwise pollen tubes would not be capable to grow significant distances before burst.

In conclusion, we have reported that the autocrine pollen tube signaling pathway regulating cell wall integrity during pollen tube growth involving Clade IB RALFs interacting with CrRLK1L-LLG receptor complexes, and with LRX/PEX cell wall proteins is partly conserved between Arabidopsis and maize, and thus likely also in other angiosperms. In contrast to Arabidopsis, pollen germination is not depending on the corresponding maize RALF homologs. We further suggest a dosage-dependent RALF signaling model integrating RALF interaction with receptor complexes and cell wall proteins, respectively, suggesting that Clade IB RALFs serve as sensors of cell wall integrity and thickness during pollen tube growth. The idea that RALFs somehow serve as sensors for cell wall integrity is not new and has been proposed before (Gonneau et al., 2018; Ge et al., 2019b), but the model presented here provides a mechanistic explanation for the burst phenotype and indirectly links the cell wall to cell surface receptors via different RALF binding affinities. Moreover, we link the regulation of pollen cell wall structure via distribution pattern of soft methyl-esterified versus stiffer de-esterified pectins and phosphorylation status of ZmPEX cell wall proteins to RALF signaling in the sub-apical region of the growing pollen tube. So far, it has only been described that FER signaling is crucial for maintaining de-esterified pectin at the filiform apparatus of the embryo sac in Arabidopsis (Duan et al., 2020). In summary, we think that the comparison of RALF signaling between slow germinating and growing pollen tubes in an eudicot species (Arabidopsis) with a fast germinating and growing monocot and grass species (maize) provided important knowledge about similarities and differences among the two species and lead to a larger number of suggestions for future research directions. We think it will now be important to better elucidate downstream signaling processes and to figure out functions of more recently evolved Clade III RALFs like ZmRALF1/5 that do not regulate cell wall integrity during pollen tube growth.

## METHODS

### Plant Materials and Transgene Transmission Studies

All maize plants were grown under the same conditions in a greenhouse at 26°C under illumination of 400 μm s^-1^ m^-2^ (measured 70 cm from the soil surface) using 400 W SON-T Agro and HPI-T Plus lamps with 16 hours light and eight hours darkness at 21°C. Air humidity was kept constant at about 60%. Pollen used for *in vitro* germination assays was collected as follows: old pollen grains were removed in the early morning by shaking and newly shed pollen grains were collected two hours later and being tested for successful germination *in vitro*. Only pollen germinating with >80% frequency was used for further analysis. Samplings were repeated at least three times within different days. For transgene transmission efficiency studies, ears were covered with white paper bags before silk emerge to avoid pollen contamination. Transgenic plants were identified by PCR and DNA agarose gel electrophoresis using RNAi-*ZmRALFs* specific primers (see details below and Table S1 for primer sequences). Transgenic/mutant lines were crossed with inbred line B73 that was used as a WT control and transmission of transgenes was detected as described above. *Arabidopsis thaliana* ecotype Columbia-0 (Col-0) seeds were sown on soil and kept for 2-3 d at 4°C in the dark for stratification. Seedlings were then grown at long-day conditions at 21°C for 16 hours of light under illumination of 150 μm s^-1^ m^-2^ and 8 hours of darkness at 18°C at 60% air humidity.

### Gene identification

To identify the *RALF* gene family in maize (*ZmRALFs*), searches with two databases were performed. ZmRALF1/2/3 mature peptide were used as queries to run BLASTP in the non-redundant protein sequences (nr) of *Zea mays* (taxi:4577) at NCBI (https://blast.ncbi.nlm.nih.gov). To avoid missing of not annotated genes, the same queries were used to run TBLASTN using the Gramene database (http://ensembl.gramene.org/). The BLOSUM62 matrix was used in both searches. Output sequences were manually selected taking into consideration a query cover >50 % and an E-score < 0.05. Selected peptides were further confirmed according to the cysteine residues arrange model (Silverstein et al., 2007). After RALF peptide identification, corresponding locus tags (B73 RefGen_v4) and gene symbols were extracted from protein introduction pages. Similar procedures were performed to obtain *ZmCrRLK1L, ZmLLG* and *ZmLRX/ZmPEX* gene families in maize. Arabidopsis FER, ANXs, BUPSs and LORELEI proteins were used as queries to BLAST the maize genome. Because of the highly variable proline-rich extension-like domain in LRX proteins (Herger et al., 2019), only ectodomains of ZmPEX1 and ZmPEX2 were used to identify the whole *ZmLRX/ZmPEX* family in the maize genome. Outputs were manually checked according to structure features (Boisson-Dernier et al., 2011; Liu et al., 2016; Herger et al., 2019).

### Plasmid Construction and Plant Transformation

Considering that nine *RALF* genes are expressed in mature maize pollen (Supplemental Figure S1), an RNAi construct was generated to downregulate the whole gene family. Gene-specific transcribed regions of *ZmRALF1/2/3* together generating about 90% *RALF* transcripts in maize pollen were cloned after DNA amplification from genomic DNA in one vector and the *RNAi-ZmRALFs* vector thereafter generated as follows: 604 bp of *ZmRALF1/2/3* including 190 bp of *ZmRALF1*, 212 bp of *ZmRALF2*, 202 bp of *ZmRALF3* that was synthesized by ThermoFisher Scientific; for primer sequences see Supplemental Table S1. After digestion with respective restriction enzymes, fragments were cloned into the corresponding splicing sites of the pUbi-iF2 vector (DNA Cloning Service) that contains the maize *Ubi* promoter. Corresponding vector regions were always sequenced. To generate CRISPR-Cas9 mutant maize plants, guide RNAs (gRNAs) were designed with Breaking-Cas (Oliveros et al., 2016) and CRISPR-Pv2.0 (Lei et al., 2014) online platforms. Corresponding sequences are included in Supplemental Table S1. The integration of gRNAs to pGW-Cas9 was carried out as previously reported (Char et al., 2017). Transformation constructs were verified by sequencing and restriction enzyme digestion. The maize inbred line HiAB hybrid was used for stable maize transformation. For *ZmRALFs* sub-localization and complementation studies in Arabidopsis, 2286 bp upstream of the ATG start codon of the Arabidopsis *AtRALF4* gene was used as endogenous promoter, followed each with ORFs of *ZmRALFs* genes cloned from maize pollen cDNA. Firstly, the *AtRALF4* promoter region was cloned from Arabidopsis genomic DNA and the promoter was integrated into pB7FWG2.0 or pB2GW7 by restriction enzyme digestion and T4 DNA ligation. Next, *ZmRALFs* fragments were transferred individually by LR reaction using Gateway LR Clonase II Enzyme Mix (ThermoFisher Scientific). Constructs containing an eGFP sequence fused C-terminally to *ZmRALFs* were cloned into pB7FWG2.0 and those lacking a tag into pB2GW7, respectively. Finally, gene constructs were transformed into *A. thaliana* ecotype Col-0.

To generate recombinant proteins in *E. coli*, predicted mature peptide regions of *ZmRALF1/2/3/5* genes were cloned into plasmid pET32b (Novagen) with gene specific primers as indicated (Supplemental Table S1). N-terminal His tag and TrxA tag were used as they were reported to promote protein solubility (Costa et al., 2014). All receptor-like kinases in this study were cloned individually into pMAL-p2p plasmid (provided by Kamila Kalinowska) with a N-terminal MBP tag, including the ectodomains of ZmFERL1/4/7/9 and ZmPEX2/4, as well as whole proteins without a signal peptide of ZmLLG1/2 and AtLRX9/10/11. Besides, ectodomains of ZmFERL7/9 were cloned into pGEX-6P1 (GE Healthcare) with a N-terminal GST tag. Constructs were transformed into *E. coli* (strain BL21).

### RNA Extraction, RT-qPCR and Gene Expression Analysis

The Trizol™ Plus RNA purification kit (ThermoFisher Scientific) was used for total RNA extraction from maize pollen. Purified RNA was reverse transcribed to cDNA using RevertAid H minus reverse transcriptase (ThermoFisher Scientific). *ZmRALFs* gene-specific quantitative PCR (qPCR) primers (Supplemental Table S1) were designed and tested in standard PCR reactions. In general, assay involved two biological replicates, each represented by two cDNA pools that were used as templates for three technical replicate reactions. Normalization was based on two internal maize reference genes, the maize membrane protein PB1A10.07c (reference gene 1, Zm00001d018359) and cullin (reference gene 2, Zm00001d024855) that were previously reported to be stably expressed (Manoli et al., 2012). Resulting calibrated normalized relative quantities were exported into Excel sheets and the ΔΔC_T_ method (Livak and Schmittgen, 2001) was used for further analysis and calculation of fold changes. To compare tissue-specific gene expression pattern of maize and Arabidopsis genes described in this study, mRNA expression levels in pollen, stigma, root, leaf and seed of maize were extracted from the CoNekT online database (https://evorepro.sbs.ntu.edu.sg/; (Julca et al., 2021). Arabidopsis pollen mRNA expression level were extracted from (Loraine et al., 2013) and expression of other tissues were obtained from Genevestigator (www.genevestigator.com). Please note that provided gene expression values of maize and Arabidopsis genes in Supplemental Figures S4 and S8 are not comparable.

### Protein Sequences and Phylogenetics Analysis

Protein sequences were download from the Universal Protein Resource (UniProt). The MUSCLE method was used for multiple sequence alignment (Edgar, 2004). The phylogenetic tree was constructed based on the Neighbor-joining statistical method, 1000 bootstrap and Dayhoff model. The phylogenetic trees were further optimized on the website of Evolview (Subramanian et al., 2019). For alignment of RALFs, predicted signal peptide and propeptide (if exists) were cut off and only predicted mature RALF sequences were used. Similarly, for alignment of CrRLK1Ls only the ectodomain were used for analysis. Proteins without signal peptide were used to align LORELEI-like GPI-anchored proteins. Sequence alignment results were shown by using Jalview (Waterhouse et al., 2009) and colored by Clustal_X (Thompson et al., 1997).

### Protein Purification and Pull-Down Assays

Purification of recombinant proteins generated in *E. coli* was modified from the protein purification protocol of The QIAexpressionist (QIAGEN) as follows: 2 mL overnight *E. coli* strains containing the corresponding protein expression plasmid were each added to 1L LB-medium with corresponding antibiotic and being incubated in a 37°C shaker at 200 rpm. Once an OD_600_ of 0.5-0.7 was reached, Isopropyl β-D-1-thiogalactopyranoside (IPTG, SERVA) was added (1 mM IPTG was used for His-tagged proteins and 0.1-0.5 mM IPTG for GST/MBP-tagged proteins) and further incubated in a 30°C shaker at 200 rpm for another 3~4 hours. Afterwards, bacteria were pelleted by centrifugation. The pellet was resuspended in 40 mL chilled lysis buffer (*His lysis buffer*: 50 mM NaH_2_PO_4_, 300 mM NaCl, 10 mM imidazole, adjust pH to 8.0; *GST/MBP lysis buffer*: 50 mM Tris pH7.5, 150 mM NaCl, 0.05% CA630 from Sigma-Aldrich) containing protease inhibitor, 0.5 mg/mL lysozyme and 1 mM PMSF. Cells were shattered by repeated sonication until samples became transparent. Lysates were collected by centrifugation at 4°C for 20 minutes at 16,000 g. Supernatants were separated from lysates and incubated with affinity beads at 4°C overnight (*His beads:* Ni-NTA from QIAGEN; *GST beads:* glutathione-agarose beads from ROTH; *MBP beads:* amylose beads from NEB). Protein-binding beads were collected by centrifugation and washed several times with washing buffer (*His washing buffer*: 50 mM NaH_2_PO_4_, 300 mM NaCl, 20 mM imidazole, adjust pH to 8.0; *GST/MBP washing buffer*: 100 mM Tris at pH 8.0) to remove unspecific binding proteins. Proteins were eluted from beads by incubating with elution buffer at 4°C for 2~4 hours (*His elution buffer*: 50 mM NaH_2_PO_4_, 300 mM NaCl, 250 mM imidazole, adjust pH to 8.0; *GST elution buffer*: 15 mg/ml reduced glutathione in 100mM Tris; *MBP elution buffer*: 25 mg/ml maltose in 100mM Tris). Proteins were further desalted with PD 10 desalting columns from GE-Healthcare. FLAG-tagged ectodomains of Arabidopsis ANX1/2 and BUPS1/2 proteins were obtained from a previous study (Ge et al., 2017). *In vitro* binding or pull-down assays were carried out as described (Ge et al., 2017). ZmRALFs and AtRALFs were generated containing a His-tag and MBP/GST-tags were used for ZmFERLs, AtANXs, AtBUPSs, ZmLLGs, ZmPEXs and AtLRXs. Briefly, samples were mixed in 500 μL binding buffer (20 mM Tris, pH7.5, 1% CA630) and rotated at 4°C for 2 hours before addition of His-beads (Ni-NTA, QIAGEN). After continuous rotating for another 2 hours, beads were pelleted and washed for five times. Samples were boiled in SDS-loading buffer and analyzed by SDS-PAGE and Western blot. For antibodies we used a His antibody (6x-His Tag Monoclonal Antibody, ThermoFisher), a MBP antibody (Anti-MBP Monoclonal Antibody, HRP conjugated, NEB), a GST antibody (GST Tag Monoclonal Antibody, ThermoFisher) and a FLAG antibody (anti-Flag-HRP, Sigma-Aldrich). For interaction studies of biotinylated AtRALF4/19 with GST-tagged ZmFERL7/9 and MBP-tagged ZmFERL1/4, the procedure was the same as described above despite that His-beads were substituted by Streptavidin Magnetic Particles (Spherotech).

### *In Vitro* Pollen Germination

Fresh maize pollen was germinated on the surface of solid pollen germination medium (PGM) in Petri dishes. First, PGM (10% sucrose (Roth), 0.005% H_3_BO_3_ (Sigma-Aldrich), 10 mM CaCl_2_ (Sigma-Aldrich), 0.05 mM KH_2_PO_4_ (Merck), 6% PEG 4000 (Merck-Schuchardt), pH 5.0) was prepared as described (Schreiber et al., 2004). After all components were mixed, an equal volume of heated 0.6% NuSieve™ GTG™ Agarose (Lonza) was added. The mixture was poured into 3 cm Petri dishes, which were ready for usage after cooling down to room temperature. Fresh pollen was obtained as described above and spread directly to Petri dishes. Pollen germination and growth status were observed and photographed by using a Nikon microscope (Nikon Eclipse TE2000-S) with a 4x objective (Plan Fluor DL 4x/0.13, PHL). Pollen germination status was divided in four categories: (i) germinated pollen growing normally, (ii) un-germinated pollen, (iii) burst pollen at germination and (iv) burst pollen tube. Calculation and statistics were made by ggpubr package in R (https://CRAN.R-project.org/package=ggpubr). *In vitro* germination of Arabidopsis pollen was performed as follows: mature pollen was germinated on the surface of solid PGM (18% sucrose, 0.01% H_3_BO_3_, 5 mM CaCl_2_, 5 mM KCl, 1 mM MgSO_4_ (Merck), pH 7.5; 1.5% NuSieve™ GTG™ agarose) in Petri dishes that were placed in humid boxes and incubated at room temperature for 4 hours. Solid PGM containing pollen tubes were cut out and placed on a microscope slide with 150 μL liquid PGM solution (lacking agarose). To visualize the cell wall, propidium iodide (PI; Sigma-Aldrich) staining was carried out. 10 mM PI in PGM solution were applied to the surface of germinated pollen tubes and incubated for 5 min. Fluorescence images were observed and collected using a ZEISS LSM 980 Airyscan2 Confocal Laser Scanning microscope with a 63x (Plan-Apochromat 63x/1.40 Oil DIC M27) objective.

### MST Assay

To test the binding affinity of maize RALF peptides with the corresponding interactors, microscale thermophoresis (MST) assays were carried out by using a Monolish NT.115 (NanoTemper Technologies). His-tagged RALF proteins were labeled with red dyes with the His-Tag labelling kit RED-tris-NTA 2nd Generation (NanoTemper) according to the manufacturers’ recommendation. His-tagged RALF proteins were diluted to 200 nM in PBST buffer (137 mM NaCl, 2.5 mM KCl, 10 mM Na_2_HPO_4_, 2 mM KH_2_PO_4_, 0.05% Tween-20 at pH 7.4). A volume of 100 μL protein solution (200 nM) was mixed with 100 μL dye solution (100 nM) and incubated at room temperature for 30 min. After centrifugation, 20 nM dye-labelled proteins were mixed with a serial dilution of interactors (ZmFERL1/4/7/9, ZmPEX2/4). Samples were loaded into glass capillaries (NT.115 Standard Treated Capillaries) and measured at 40% MST power and 10% LED power. Recorded data were analyzed with MO.Affinity Analysis Software v2.3 (NanoTemper Technologies).

### Phospho-Proteome Analysis

To compare the phospho-proteome of wild type (B73) and RNAi-Zm*RALFs* mutant lines, fresh pollen was germinated *in vitro* for 45 minutes on solid PGM as described above. Successful pollen germination and tube formation was checked by using an inverted microscope (Nikon Eclipse TE2000-S) with a 4x objective (Plan Fluor DL 4x/0.13, PHL). Pollen tubes were filtered by PluriStrainer^®^ 40 μm (pluriSelect) and washed twice with fresh PGM. 50 mg of pollen tube samples were transferred from the surface of the strainer into a 1.5 mL collection tube and immediately frozen in liquid nitrogen and stored at −80 °C for phospho-proteome analysis. The phospho-proteome was generated as described (Mergner et al., 2020). 100 μg precipitated, washed and in urea buffer re-suspended total protein was digested with trypsin. TMT labeling, phosphopeptide enrichment and fractionation was performed as described previously (Ruprecht et al., 2017; Zecha et al., 2019). Nano-flow LC-ESI-MS measurements were performed with a SPS-MS3 method on a Fusion Lumos Tribrid mass spectrometer (Thermo Fisher Scientific) coupled to a Dionex 3000 HPLC (Thermo Fisher Scientific). Full-scan MS1 spectra (m/z 360-1300) were acquired with 60,000 resolution, an automatic gain control (AGC) target value of 4e5 and 50 ms maximum injection time. Precursor ions were fragmented via CID (NCE of 35%, activation Q of 0.25) and recorded in the Orbitrap at 15,000 resolution (isolation window 0.7 m/z, AGC target value of 5e4, maxIT of 22 ms). The 10 most intense peptide fragments for each precursor were selected in the ion trap, further fragmented via HCD (NCE of 55%) and read out in the Orbitrap for MS3 quantification (50,000 resolution, charge dependent isolation window from 1.2 m/z (2+) to 0.7 (5-6+), AGC of 1.2e5, maxIT of 120 ms). Peptide and protein identification and quantification was performed with MaxQuant (Cox and Mann, 2008) using standard settings (version 1.6.0.16).

### Immunohistochemistry

For fluorescence labelling of maize pollen tube cell wall epitopes, fresh pollen grains were collected, and germinated in liquid PGM within ibiTreat μ-Slide VI 0.4 channels (Ibidi). 45 min after germination, liquid PGM was replaced with fixative solution (100 mM PIPES buffer, pH 6.9, 4 mM MgSO_4_, 4 mM EGTA, 10% (w/v) sucrose, and 5% (w/v) formaldehyde) and pollen tubes were fixed at room temperature for 90 min. After washing with PBS buffer (137 mM NaCl, 2.5 mM KCl, 10 mM Na_2_HPO_4_, 2 mM KH_2_PO_4_ at pH 7.4), pollen tubes were incubated with PBS buffer containing 3.5% (w/v) BSA at room temperature for 30 min and then being washed by PBS buffer. All antibodies were diluted in PBS buffer with 3.5% (w/v) BSA. Incubation was done at 4 °C overnight for the primary antibody and 3 hours at 30 °C for the secondary antibody. De-esterified and esterified pectins were labelled with JIM5 and LM20 antibodies, respectively (diluted 1:50, Kerafast) followed by Alexa Fluor 594 anti-rat IgG (diluted 1:100, Thermo Fisher Scientific) staining. Negative controls were performed by omitting primary or secondary antibody incubation. Pollen tubes were finally kept in ROTI^®^Mount FluorCare (Carl Roth) and fluorescence images were collected using a ZEISS LSM 980 Airyscan2 Confocal Laser Scanning microscope with a 63x (Plan-Apochromat 63x/1.40 Oil DIC M27) objective.

### Accession numbers

Accession numbers of experimentally analyzed genes in this study are as follows: *ZmRALF1* (Zm00001d039766), *ZmRALF2* (Zm00001d008881), *ZmRALF3* (Zm00001d039429) *ZmRALF5* (Zm00001d019239)*, ZmFERL1* (Zm00001d029047), *ZmFERL4* (Zm00001d037839), *ZmFERL7* (Zm00001d035992), *ZmFERL9* (Zm00001d045515), *ZmLLG1* (Zm00001d003139), *ZmLLG2* (Zm00001d020876), *ZmLLG3* (Zm00001d051672), *ZmLLG4* (Zm00001d017874), *ZmPEX1* (LOC103648188), *ZmPEX2* (Zm00001d048715), *ZmPEX3* (Zm00001d023858), *ZmPEX4* (Zm00001d040802), *AtRALF4* (At1G28270), *AtRALF19* (At2G33775), *ANX1* (At3G04690), *ANX2* (At5G28680), *BUPS1* (At4G39110), *BUPS2* (At2G21480), *AtLRX9* (At1G49490), *AtLRX10* (At2G15880), *AtLRX11* (At4G33970). See Supplemental Table S2 for all genes.

## SUPPLEMENTAL MATERIAL

**Supplemental Figure 1.** Expression pattern of 24 maize *RALF* genes in indicated tissues.

**Supplemental Figure 2.** RALF protein sequences from maize and Arabidopsis diverge across three major phylogenetic clades.

**Supplemental Figure 3.** Relative expression levels of three *ZmRALFs* in pollen tubes of WT compared with six *RNAi-ZmRALFs* mutant lines.

**Supplemental Figure 4.** Phylogenetic and expression analysis of 17 maize CrRLKs1L receptor kinases to identify putative orthologs among 17 Arabidopsis CrRLK1Ls.

**Supplemental Figure 5.** Phylogenetic tree of 34 CrRLK1L proteins from maize and Arabidopsis showing their domain organization and expression pattern patterns.

**Supplemental Figure 6.** Maize CrRLK1L receptor kinases are capable to interact with Clade IB Arabidopsis RALFs and *vice versa*.

**Supplemental Figure 7.** ZmPEX2 phosphorylation status in pollen tubes of WT and two RNAi-Zm*RALFs* mutant lines as indicated.

**Supplemental Figure 8.** Phylogenetic analysis and expression pattern of PEX/LRX proteins and genes from maize and Arabidopsis.

**Supplemental Figure 9.** Sequence alignment of pollen-expressed LRR-extensin-like cell wall proteins named as PEX in maize and LRX in Arabidopsis, and their phosphorylation sites.

**Supplemental Figure 10.** Maize Clade IB RALFs are capable to interact with Arabidopsis LRX proteins.

**Supplemental Figure 11.** Disturbance of cell wall organization in *ZmRALFs* mutants.

**Supplemental Table 1.** List of PCR primers used in this study.

**Supplemental Table 2.** Gene ID and protein information of all maize ZmRALFs, ZmFERLs, ZmPEX/LRXs and ZmLLGs.

## ACKNOWLEDGMENTS

We thank Kamila Kalinowska (University of Regensburg) for providing the pMAL-p2p vector and Armin Hildebrand for plant care. The German Research Foundation (DFG) is acknowledged for financial support via SFB924 (to TD and BK) and SFB960 (to GL), and the China Scholarship Council (CSC) for a fellowship to LW.

## AUTHOR CONTRIBUTIONS

TD designed the project and LZZ together with LW performed most of the experiments. TD, LZZ, GL and LJQ supervised the study and contributed with analyses and data interpretation. ZG and XL contributed with material and analyses, while JM and BK generated and analyzed phospho-proteomic data. LZZ, LW and TD wrote the article with input from all authors.

